# Nano-flow cytometry of single extracellular vesicles reveals subpopulation differences across cell types and pharmacological perturbations

**DOI:** 10.1101/2025.07.09.663918

**Authors:** Nathalie Nevo, Alix Zhou, Nicolas Ansart, Lea Cohen Attali, Eric Rubinstein, Coralie Guérin, Lorena Martin-Jaular, Clotilde Théry

## Abstract

Extracellular vesicles (EVs) are lipid bilayer-enclosed particles released by most cell types, which can transfer signals and cargoes between cells. EVs released by a single donor cell source are increasingly recognized as extremely heterogeneous, in terms of size, intracellular origin, and cargo composition. Analyzing large numbers of EVs at the single vesicle level is therefore the only way to truly decipher their heterogeneity. Here, we developed a reliable pipeline of single EV analysis using a nanoparticle-dedicated flow cytometer, which detects particles and measures their size down to 55 nm in diameter, without the need for vesicle pre-immobilization or fluorescent label. We show that titrating each antibody, eliminating unbound antibodies and using EVs devoid of the analyzed markers as negative controls are required to reliably quantify the proportion of EVs bearing none or any combination of two markers, as well as to measure their sizes. We thus observed, depending on the cell source (human cell lines MDA-MB-231, HeLa, A549), variable proportions of EVs bearing none of the CD9, CD81 and CD63 tetraspanins often used to define EVs, and of single- and double-positive EVs for each of these markers. We also observed CD29 (ITGB1) as a protein detected as frequently on EVs as CD9, while other transmembrane proteins (CD44, SSEA-4, CD98), were detected in a small proportion of EVs, and mostly of relatively large size. Finally, we used this pipeline to uncover differential effects of small molecule drugs on subtypes of EVs, and showed that Homosalate increased the proportion of CD9+/CD63+ EVs while two other drugs, Dipivefrin hydrochloride and Metaraminol bitartrate, instead increased the proportion of CD9-/CD63+ EVs. Overall, nano-flow cytometry allows to reliably quantify proportions of EV subpopulations suggested by bulk analyzes of EV markers, at single EV resolution.

## INTRODUCTION

Extracellular vesicles (EVs) are lipid bilayer-enclosed particles released by most cell types (Cocozza *et al*, 2020a). EVs are able to transfer signals and cargoes between donor and recipient cells and are thus instrumental in intercellular communication (Yates *et al*, 2022a, b). However, EVs released by a single donor cell source are increasingly recognized as extremely heterogeneous, in terms of size, intracellular origin, and cargo composition (Dixson *et al*, 2023; van Niel *et al*, 2022). For instance, EVs can be formed by direct outward budding from the plasma membrane, leading to the release of ectosomes (also called microvesicles). Alternatively, EVs can result from initial inward budding into intracellular compartments of the endocytic pathway, thus generating intraluminal vesicles of multivesicular compartments/bodies (MVBs), which will become exosomes, when released upon fusion of MVBs with the plasma membrane. Since other budding events at the plasma membrane (for instance, from the base or the tip of plasma membrane extensions) or other intracellular compartments (for instance, autophagosomes) can also be sources of EVs (Buzas, 2023; Dixson *et al*., 2023), the EV preparations used for therapeutic applications or for basic science studies always contain a heterogeneous mixture of different vesicles. EV characterization by the presence of given proteins in bulk preparations therefore does not give accurate information on the nature and proportion of the different EV subtypes present in the mixed population.

Analyzing EVs at the single vesicle level is the only way to decipher their heterogeneity, but it is made difficult by their small size, often below the resolution threshold of conventional fluorescence microscopes and flow cytometers. Flow cytometry, by analyzing particles in suspension and passing one-by-one under laser illumination, seems the optimal approach for unbiased analysis of large enough numbers of single EVs for reliable quantification. It avoids the need for pre-immobilization on a rigid surface required for fluorescence or super-resolution-based microscopy, which is always susceptible to exclude particles that could not be captured.

Technical improvements in the last decades have allowed the development of flow cytometry instruments able to analyze particles smaller than about 200 nm in diameter, which would fall into the background noise of classical flow cytometers. First uses for single EV analysis of in-house improved instruments were described about a decade ago (Nolte-’t Hoen *et al*, 2012; Zhu *et al*, 2014), but they required a strong expertise in flow cytometry, and were thus difficult to implement in other laboratories. In the last 5 years, commercial flow cytometers dedicated to nanoparticles have been released, with the ability to detect particles smaller than 300 nm in diameter based on their size scatter, independently of any pre-labeling by fluorescent dyes, thus avoiding the bias induced by potential non-universal labeling of particles. EVs have a lower refractive index as metal or polymer beads, but they can still be detected down to 40-50 nanometers of diameter, and fluorescence can then be measured on each individual particle after labeling with fluorescent antibodies or dyes (Brealey *et al*, 2024; Chen *et al*, 2023; Tian *et al*, 2020).

Here, we aimed to use one of the commercial nanoparticle-dedicated flow cytometers, to quantify the proportions of EVs bearing different combinations of surface markers, from different sources of human cell lines, and their changes upon cells’ treatment by small molecule drugs we had used previously to modify EV release (Grisard *et al*, 2022). The three tetraspanins CD9, CD63 and CD81 are often used as universal markers of small EVs, however, recent cell biology and differential proteomic studies from us and others (Martin-Jaular *et al*, 2021; Mathieu *et al*, 2021; Palmulli *et al*, 2024) suggest that they are probably differently enriched in exosomes (mostly CD63) versus ectosomes (mostly CD9, maybe CD81). We thus first focused on these molecules to establish a robust protocol for specific EV staining, using EVs from cells knocked out for each of them as negative controls. We could then implement the labeling and analysis protocol 1) on EVs from different human cell sources (tumor cell lines HeLa, MDA-MB-231, A549), 2) using antibodies to other surface molecules (ITGB1, CD44, SSEA4, CD98), 3) on EVs produced under different environmental conditions of the source cells (treatment with off-patent small molecule drugs: Homosalate, Dipivefrin hydrochloride, Metaraminol bitartrate and Ebselen). Our results demonstrate clear differences in proportions of tetraspanin-negative, CD63, CD9 and CD81 single- or double-positive EVs in these different situations, and pave the way to identify drugs able to specifically modulate secretion of a given subtype of EVs.

## RESULTS

### Establishment of the conditions for specific staining of EVs with antibodies for nano-flow cytometry

Throughout this study, we used a flow cytometer specifically designed to analyze small particles of sub-micron size for side scatter and 2 fluorescent signals simultaneously (U30 Flow NanoAnalyzer, NanoFCM Inc.). Our first aim was to establish a standardized and reproducible protocol to allow conclusive identification of antibody-labeled EVs among a bulk preparation. To do so, we followed recommendations of ISEV for EV cytometry, especially for the importance of negative and procedural controls (Welsh *et al*, 2020). We used EVs from HeLa WT cells and HeLa CD81 knockout (KO) cells as a negative control. EVs were isolated by size exclusion chromatography (SEC) from conditioned culture media (CM) and subsequently labeled with anti-CD81 APC antibody (clone 5A6) in a serial dilution range from 1 µg/ml to 20 µg/ml. We first, as suggested by the instrument provider, ran EVs in the Flow NanoAnalyzer after diluting the samples 1:100 in PBS without any additional cleaning step (Fig. 1A). We observed that the percentage of CD81-positive EVs increased with antibody concentration for both WT and CD81 KO EVs, with minor differences between the positive and negative EVs at most concentrations of antibodies, especially 47% positive CD81-KO EVs when using 4 µg/ml of antibody. We confirmed by imaging flow cytometry of the cells, that HeLa CD81 KO cells were negative for CD81 (Sup. Fig. 1), ruling out that EVs isolated from these cells could bear CD81. Therefore, PBS dilution did not eliminate detection of unbound or non-specifically bound antibodies. We next passed the antibody-labeled EVs through small SEC columns packed with beads retaining particles smaller than around 30 nm in diameter, to separate EVs from small soluble components such as free antibodies, before running them in the Flow NanoAnalyzer. The percentage of CD81-positive EVs also increased with antibody concentration, but at antibody concentrations of 4 µg/ml or lower, CD81 KO EVs were negative for CD81 (Fig. 1B). These results demonstrate that small SEC columns are effective at removing free and non-specifically bound antibodies from the EVs. We then analyzed the percentage of EVs from HeLa or MDA-MB-231 cells bearing CD9, CD81 or CD63. For all labeling experiments presented in this study, antibodies were first titrated on human HeLa WT EVs, with HeLa KO EVs as a negative control, and/or MDA-MB-231 EVs with E0771 murine cell EVs as a negative control. Fig. 1C shows examples of the dot plots and gating established for the anti-CD9, anti-CD63 and anti-CD81 antibodies coupled to FITC or APC. To determine the optimal concentration of each antibody used, we plotted the stain index curve as a function of antibody concentration (Sup. Fig. 2), and compared this curve with the fluorescence signal observed on negative control EVs (dot plots of E0771 EVs labeled with different concentrations of anti-CD63-APC are shown in Sup. Fig. 2 as illustration). An arrow on each graph indicates the optimal concentration chosen to ensure specific labeling of EVs without labeling the corresponding negative control EVs: due to the observed signal in negative EVs, the optimal concentration could be, for some antibodies, lower than the concentration giving the highest stain index. A summary table (Table 1) outlines the important parameters we established as necessary for sensitive and specific labeling of EVs before Flow NanoAnalysis, which can be used for any protein of interest. All subsequent results presented here were obtained following this pipeline. Sup. Table 1 provides informations on each antibody tested here and the optimal concentration for EV staining (in most cases for our experimental samples: between 2 and 4 μg/ml).

**Figure 1:**
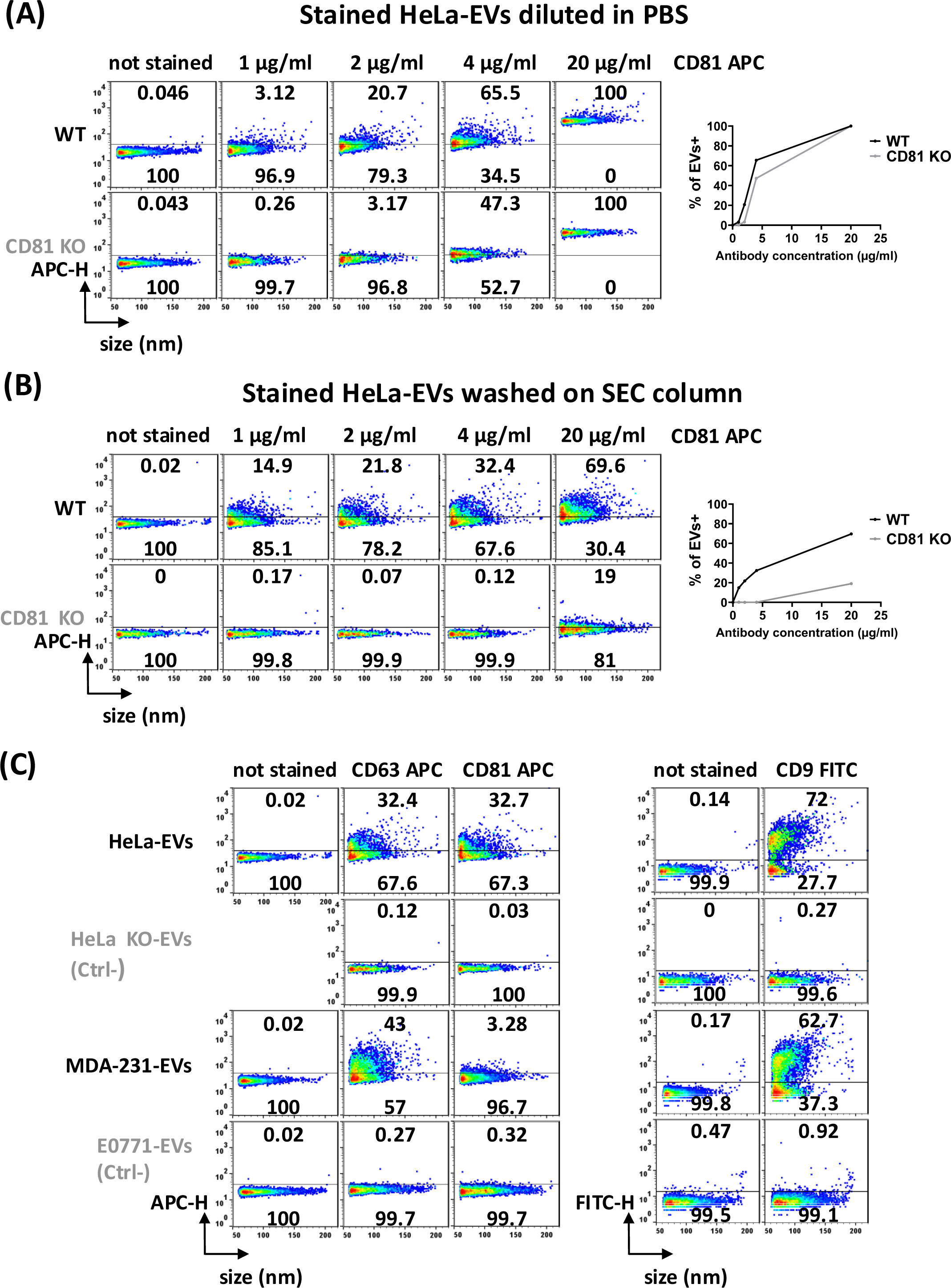
Establishment of optimal staining conditions to characterize the tetraspanin composition of EVs. A-B) Determination of staining conditions for single EV Flow NanoAnalysis using EVs from HeLa WT or CD81 KO cells. HeLa WT and HeLa CD81 KO EVs were labeled with anti-human CD81-APC antibody at concentrations ranging from 1 µg/ml to 20 µg/ml for 30 minutes at 4°C. EVs from the control condition (not stained) were treated in the same manner as the labeled EVs, except that they were not incubated with the antibody. Dot plots of particle size distribution in nm versus APC-H fluorescence intensity are shown (left panel), with the percent of EVs below or above the positivity threshold indicated. Curve of percent of EVs positive for CD81 as a function of the final concentration of antibody used for staining (right panel). A) Labeled EVs were diluted 100 times with PBS and then analyzed by Flow NanoAnalyzer. B) Labeled EVs were passed through a mini-SEC column and eluted with PBS before being analyzed by Flow NanoAnalyzer. C) Representative dot plots of EVs stained for CD63, CD81 or CD9. Representative dot plots of HeLa WT EVs versus HeLa KO EVs: for each staining CD63 APC, CD81 APC or CD9 FITC, the corresponding HeLa KO EVs was CD63 KO, CD81 KO and CD9 KO respectively. HeLa WT and HeLa KO EVs were stained with the same dilution of antibody for each condition. Representative dot plots of MDA-MB-231-EVs (MDA-231-EVs) versus mouse E0771-EVs: for each staining CD63 APC, CD81 APC or CD9 FITC, E0771 EVs were stained with the same dilution of antibody. The results of the analysis are represented by size distribution in nm and fluorescence intensity (APC-H or FITC-H), with the percent of EVs below and above the positivity threshold indicated.

**Table 1:**
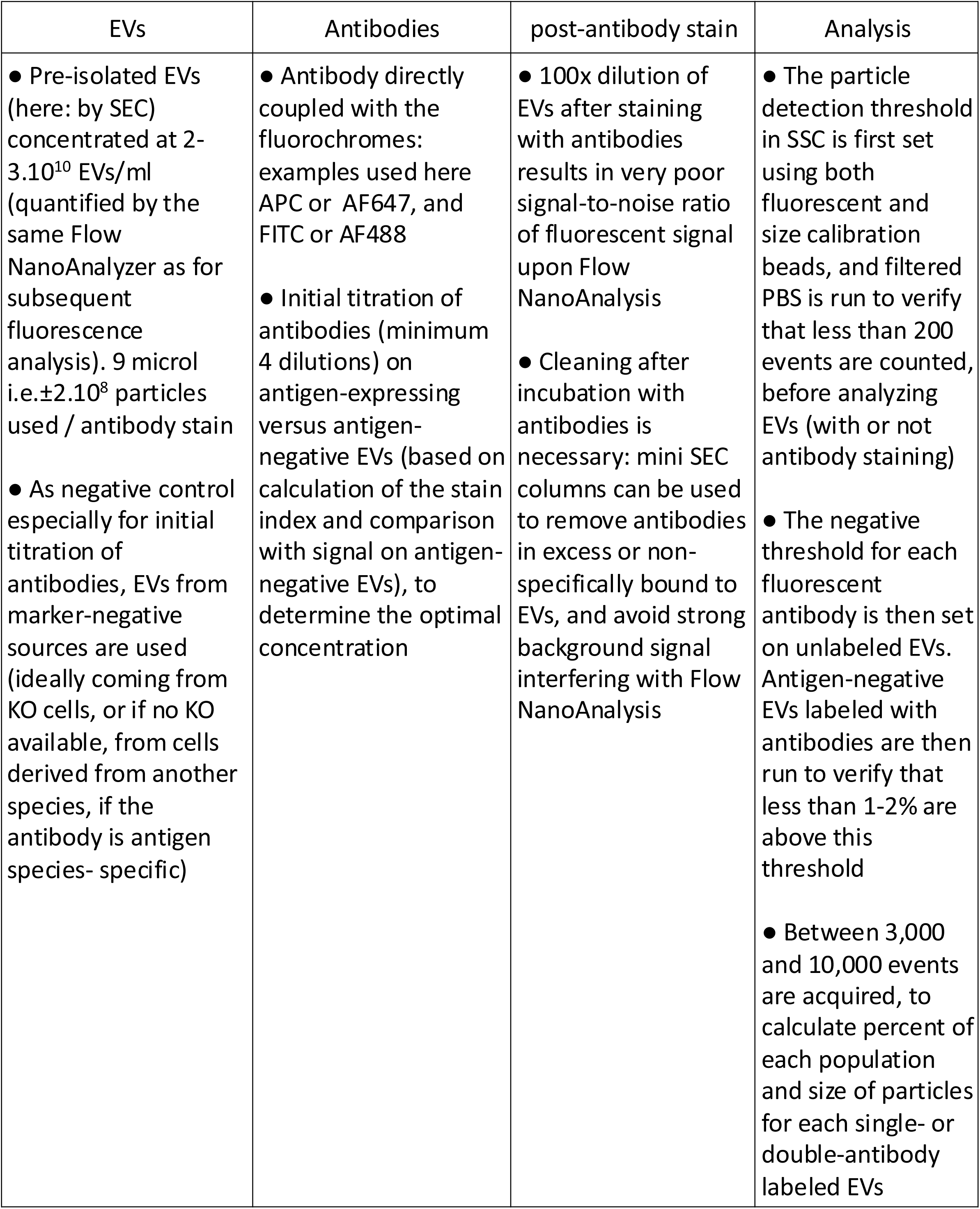
Key parameters required to achieve specific labeling and sensitive detection of EVs stained with fluorescent antibodies by Flow NanoAnalysis: 1/ EV concentration and sources 2/ Antibodies 3/ Post-staining steps 4/ Parameters of analysis.

### Proportions and size of EVs bearing the CD63, CD81 and CD9 tetraspanins vary in different cell lines

We first used our protocol for single staining with fluorescent antibodies, to characterize the EVs released by HeLa, MDA-MB-231 and A549 cells (respectively cervical, breast and lung carcinoma), in terms of percentage of EVs expressing the tetraspanins CD63, CD81, and CD9, which are often considered as “universal” markers of EVs (Fig. 2A). We could observe clear differences between these three cells in the proportion of EVs bearing each of these markers: less than 1/3 of HeLa (28.3±4.4%) or A549 (22.3±1.5%) EVs compared to almost 1/2 of MDA-MB-231 EVs (43.5±2.2%) were positive for CD63, while more than 2/3 for HeLa (72.2±4.9%) and only half or slightly more for MDA-MB-231 (57.6±5.2%) and A549 (54.8±3.5%) were positive for CD9. For CD81 the difference was even more striking, since almost half of A549 EVs (39.3±3.2%), 1/4 of HeLa EVs (27.4±3.4%) while hardly any MDA-MB-231 EVs (4.8±2.2%) were positive. MDA-MB-231 therefore releases proportionally more CD63+ EVs than HeLa and A549 cells, and hardly any CD81+ EVs, while HeLa cells release more abundant CD9+ EVs than the other two cell lines.

**Figure 2:**
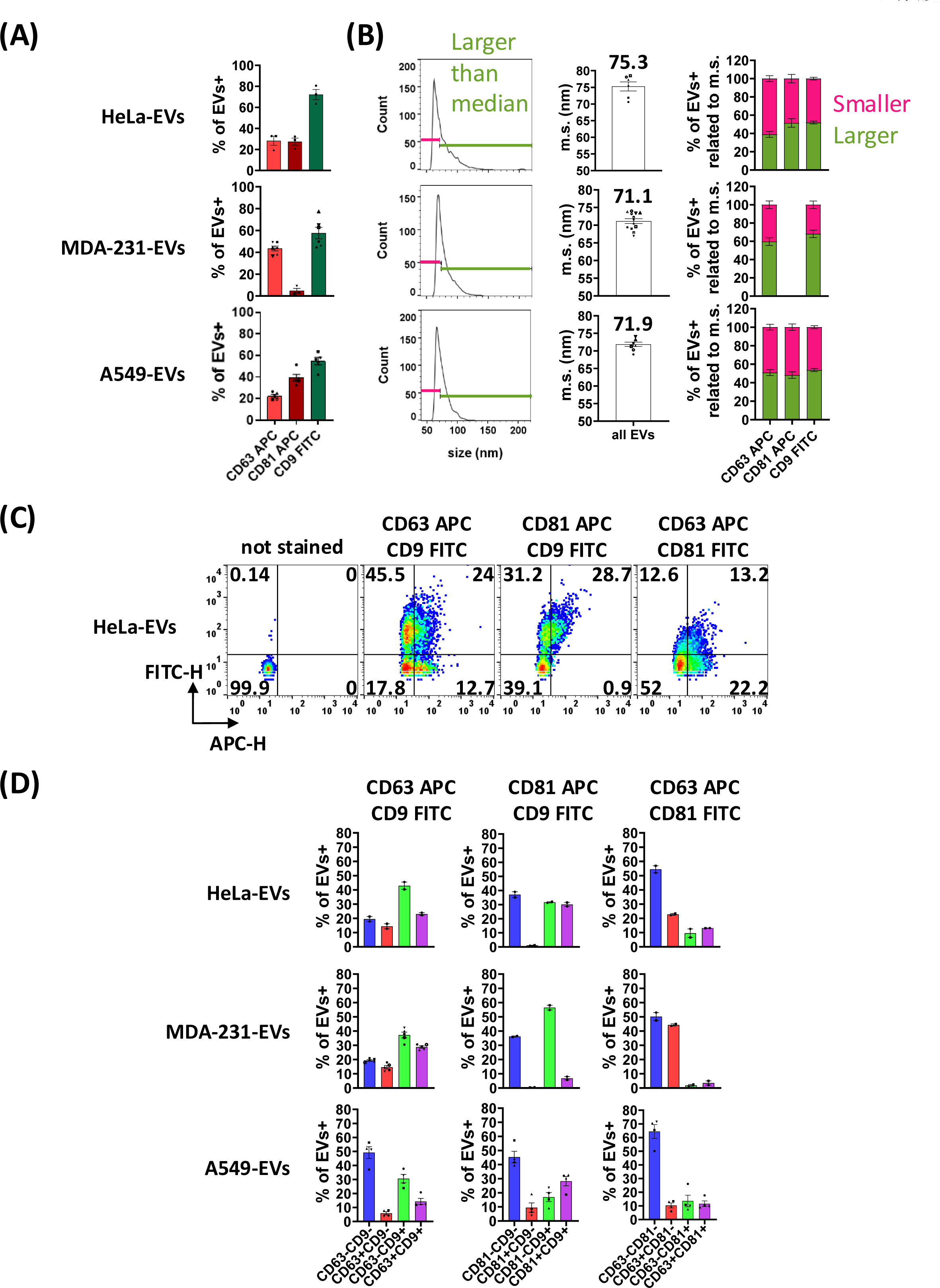
Single and double stainings of HeLa, MDA-MB-231 and A549 EVs for the tetraspanins CD63, CD81 and CD9. A-B) Single staining of Hela-, MDA-MB-231- and A549-EVs for the tetraspanins CD63, CD81 and CD9. A) Bar charts showing the percent of EVs labeled for CD63 APC, CD81 APC and CD9 FITC among the total population for each cell line, as indicated (Data are expressed as mean ± SEM, of n=3-6 independent experiments represented each by a different symbol). B) Representative size distribution profiles for HeLa-, MDA-MB-231- and A549-EVs (left panel), and median size (m.s.) in nm of the EV population calculated in 6 (HeLa), 13 (MDA-MB-231) or 8 (A549) independent non-stained EV preparations (middle panel). Mean of the median size of HeLa-, MDA-MB-231- and A549-EVs calculated by Flow NanoAnalyzer is indicated above each graph. Right panel: proportion of EVs positive for CD63-APC, CD81-APC or CD9-FITC of size above (green) or below (pink) the median size of total EVs. CD9+ and CD81+ HeLa-EV are equally distributed above and below median size, whereas CD63+ EVs are more prominently among the lower-size EVs (38% above median size). For MDA-MB-231 EVs, both CD63+ and CD9+ are more prominently in the larger than median size EVs (above 60%). For A549, CD63+, CD81+ and CD9+ EV are equally distributed above and below the median size. C-D) Double staining of Hela-, MDA-MB-231- and A549-EVs for CD63, CD81 and CD9. C) Representative dot plots APC-H versus FITC-H of EVs from HeLa cells not stained, or stained for CD63 APC/CD9 FITC, CD81 APC/CD9 FITC orCD63 APC/CD81 FITC. The position of the threshold for positive events was established on unstained EVs, and the percent of EVs in each quadrant is indicated. C) Bar charts showing the percentage of the double-negative population (blue), single-positive population (red: APC, green: FITC) and double-positive population (purple) obtained for each double staining for HeLa-, MDA-MB-231- and A549-EVs (mean ± SEM of n=2-5 independent experiments). Comparison of proportions of tetraspanin-positive EVs calculated from single versus double stainings are shown in Sup. Fig. 3C.

Of note, similar proportions of positive EVs were observed when using other clones of anti-CD81 antibodies, and/or other fluorochromes coupled to the same antibodies: Sup. Fig.3A,B show results on EVs from HeLa, A549 or MDA-MB-231 stained with the anti-CD81 5A6 clone coupled to FITC or to APC, on HeLa and MDA-MB-231 EVs stained with another anti-CD81 clone coupled to APC (clone provided by Miltenyi with the MacsPlexEV kit), and of MDA-MB-231 EVs stained with antibodies to CD63 and to CD29 (relevant to Fig.3) bound to different fluorochromes (Sup. Fig. 3B).

**Figure 3:**
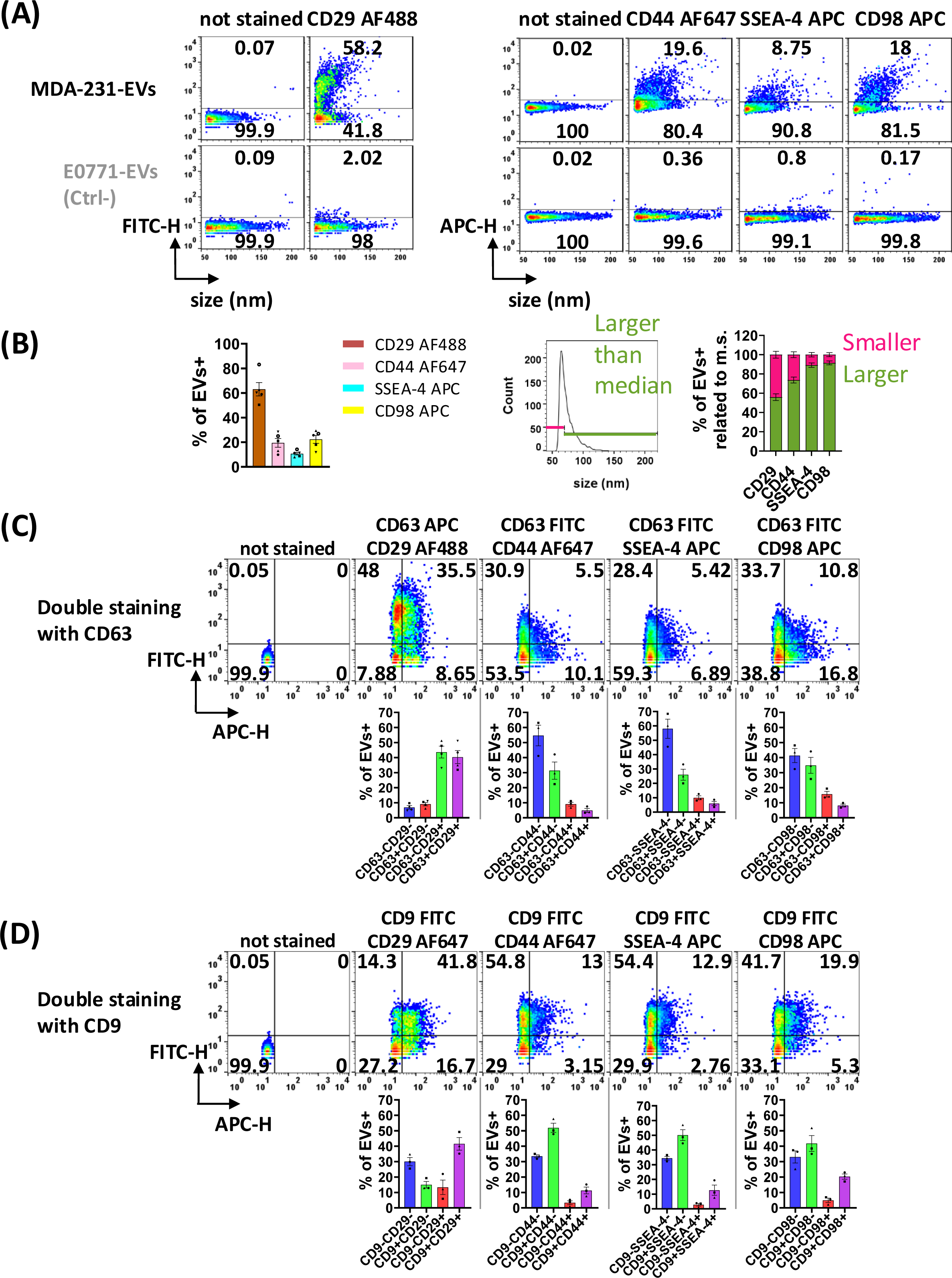
Single and double stainings of MDA-MB-231-EVs for CD29, CD44, SSEA-4 and CD98. A) Representative dot plots of single staining of MDA-MB-231-EVs versus mouse E0771-EVs (negative control) for CD29 AF488, CD44 AF647, SSEA-4 APC and CD98 APC. The results are represented with size distribution in nm and fluorescence intensity (FITC-H or APC-H), and the percentage of EVs below or above the positivity threshold is indicated on each dot plot. B) Bar charts showing the percentage of MDA-MB-231-EVs positive for CD29 AF488, CD44 AF647, SSEA-4 APC or CD98 APC compared to the total population (mean ± SEM of n=5 independent experiments) (left panel). Representative histogram (middle panel) and proportion of EVs positive for CD29, CD44, SSEA-4 or CD98 of size above (green) or below (pink) the median size of total EVs (right panel). CD29+ EVs are more prominently in the larger than median size EVs (56%), like CD9+ and CD63+ EVs (Fig. 2B). EVs positive for the other 3 markers are even more predominantly in the large EV population (74 to 91%). C-D) Representative dot plots (top panels) and quantification (bottom panels) of double staining of MDA-MB-231-EVs for one of the 4 markers (CD29, CD44, SSEA-4 or CD98) with CD63 (C), or with CD9 (D). Bar charts show the percentages of double-negative (blue), single-positive (green: AF488 or FITC, red: AF647 or APC) and double-positive (purple) EV populations of the corresponding stainings (mean ± SEM of n=3 independent experiments for all double stainings except n=4 independent experiments for CD63 APC/CD29 AF488).

Another interesting information provided by Flow NanoAnalyzer is the size of EVs: this parameter is not an absolute value, since it is calculated from their SSC signal intensity and dependent on their refractive index, but it still allows to identify different subpopulations of EVs within the bulk preparation. We thus next calculated the median size of the total EV population, and the percentage of EVs larger versus smaller than the median size among those bearing CD63, CD81 or CD9 (Fig. 2B). For HeLa EVs, the median size was 75.3±1.4 nm: only 38.9±3.2% of EVs larger than the median size were positive for CD63, while CD81+ and CD9+ EVs were evenly distributed above and below the median size (respectively above median of 51.5±4.7% for CD81, and 52±1.5% for CD9). This suggests that CD63 is more prevalent on smaller vesicles compared to CD81 and CD9. For MDA-MB-231 EVs, the median size was slightly smaller (71.1±0.7 nm), and on average, 59.7±4.2% of EVs larger than the median size were positive for CD63, as compared to 68.1±4.1% positive for CD9. The median size of A549 EVs was similar to that of MDA-MB-231 EVs (71.9±0.6 nm), but here, CD63+, CD81+ and CD9+ EVs were evenly distributed above and below the median size (respectively above median of 50.9±3.1% for CD63, 48.3±3.4% for CD81, and 53.7±1.6% for CD9) (Fig. 2B).

To better characterize the subtypes of EV populations released by HeLa, MDA-MB-231 and A549 cells, we next performed double labeling of tetraspanins on EVs. Examples of dot plots and gating for HeLa EVs are shown in Fig. 2C, and quantifications of single- and double-positive populations of EVs in Fig. 2D. As shown in Sup. Fig. 3C, for all 3 cell lines, the percentages of EVs bearing each tetraspanin were similar when calculated from the single staining (as in Fig. 2A), or from the double stainings (as in Fig. 2D) by summing up single- and double-labeled EVs, thus validating our protocol for Flow NanoAnalysis. A first observation from the double-staining was that approximately 20% of particles were neither CD63+ nor CD9+, both in HeLa and MDA-MB-231 EVs, and even more (almost 50%) in A549 EVs. Since CD81 is detected exclusively together with CD9 on HeLa and MDA-MB-231 EVs (Fig. 2D, middle panels), the 20% of CD63-/CD9-EVs from these cells were necessarily also CD81-negative. For A549, the high percentage of CD63/CD9-negative EVs was also observed upon double staining with anti-CD81/CD9 or anti-CD63/CD81, suggesting that a large proportion of A549 EVs do not bear any of these classical EV markers. For the CD63/CD9-labeled EVs, the profiles of EVs positive for each marker were similar between HeLa and MDA-MB-231, although there was a trend toward a higher proportion of CD63-CD9+ EVs and a lower proportion of CD63+CD9+ EVs in HeLa-compared to MDA-MB-231-derived EVs. The A549 EVs displayed a lower proportion of double-positive EVs than those of the other 2 cell lines. Interestingly, when performing the CD81/CD9 double labeling, we observed that all CD81+ EVs from HeLa cells were also CD9+ (1.1±0.2% of CD81+/CD9-EVs versus 30.1±1.4% of CD81+/CD9+ EVs), which was not the case for A549 EVs (9.5±3.3% of CD81+/CD9-EVs versus 28.2±3.2% of CD81+/CD9+ EVs). Finally, with the CD63/CD81 double labeling in HeLa and A549 EVs, we observed that CD63 and CD81 were only minimally co-localized on the EVs (13.1±0.05% or 11.6±2.1% CD63+CD81+, respectively).

Overall, we demonstrate here that the Flow NanoAnalyzer technology, coupled with a protocol we developed and optimized, enables precise and reproducible quantification of the percentages of the tetraspanins CD63, CD81, and CD9 present on the surface of EVs. Furthermore, we show that three different tumor cell lines release distinct EV subpopulations with different tetraspanin profiles, and different proportions of EVs bearing no detectable levels of these tetraspanins.

### Distribution and size of MDA-MB-231-EVs bearing other surface markers are similar (CD29) or different (CD44, SSEA-4, CD98) from those of CD9- or CD63-bearing EVs

In a previous work (Grisard *et al*., 2022), using a multiplex bead-based flow cytometry technique (MacsPlexEV, Miltenyi) that combines capture beads conjugated to a panel of 37 antibodies and detection by a mixture of antibodies against the tetraspanins CD63, CD81, and CD9, we had identified the proteins CD29 (integrin β1/ITGB1), CD44 (Epican), and the ganglioside SSEA-4 (glycolipid carbohydrate epitope stage-specific embryonic antigen 4) as expressed on the surface of tetraspanin-positive MDA-MB-231-EVs. CD29, CD44, and SSEA-4 were also co-detected with the protein SLC3A2/CD98, identified as an interesting EV surface protein in our previous work (Grisard *et al*., 2022; Mathieu *et al*., 2021). We confirmed the MacsPlexEV results here (Sup. Fig. 4). We thus selected antibodies specific for these 4 proteins to determine whether our Flow NanoAnalysis pipeline could also be implemented to analyze other surface markers of EVs than the 3 tetraspanins. Similar as for the tetraspanins, we established stain index curves based on antibody concentration and verified that control EVs (mouse E0771-EVs) were negative for the selected concentrations of anti-CD29 AF488 or AF647, anti-CD44 AF647, anti-SSEA-4 APC, and anti-CD98 APC (Sup. Fig. 2 and Fig. 3A). Representative dot plots (Fig. 3A) showed distinct profiles in terms of percentages of positive vesicles and their distribution according to vesicle size for these 4 antibodies. Quantitative analysis (Fig. 3B) showed CD29 presence on 63.1±5.4% of MDA-MB-231-EVs, which is comparable to the percentage of CD9+ EVs (Fig. 2A), thus consistent with the high signal obtained for CD29 in the multiplex bead-based flow assay (Sup. Fig. 4). EVs were positive to a lesser extent for CD44 (19.5±3.5%), SSEA-4 (10.7±1.1%), and CD98 (22.4±3.2%) (Fig. 3B, left). In terms of size (Fig. 3B, right), CD29+ EVs were, like CD9+ EVs (Fig. 2B), found mainly on vesicles larger than the median size (56.0±3.5%). Interestingly, the percentage of positive EVs larger than the median vesicle size was even higher for CD44, SSEA-4, and CD98 (73.7±3.1%, 89.1±2.3%, and 91.5±2.1%, respectively).

We then performed double staining of the new markers with either CD63 (Fig. 3C) or CD9 (Fig. 3D) on MDA-MB-231-EVs. We first verified that % of positive EVs were similar when using either FITC- or APC-labeled anti-CD63, and either AF488- or AF647-labeled anti-CD29 (Sup. Fig. 3B). Double labeling of CD63/CD29 (Fig. 3C) and CD63/CD9 (Fig. 2D) displayed slightly different profiles with more double-positive EVs CD63+CD29+ than CD63+CD9+ (40% vs 29%), and with more single-positive CD29+ than CD9+ EVs (44% vs 37%). Double labeling of CD9/CD29 showed the same proportion of double-positive EVs as for CD63/CD29 (41.5±4.1%) and only around 13-15% of single-positive EVs for both markers, showing that CD9 and CD29 are often present on the same EVs. For the other surface markers, EVs bearing CD63 together with CD44, SSEA-4, or CD98 were very scarce (5 to 8%), and twice less represented than those bearing these markers alone (9 to 16%), while, by contrast, EVs bearing CD9 together with CD44, SSEA-4 or CD98 were four times more abundant than those bearing each of these markers alone. In conclusion, in the population of EVs released by MDA-MB-231, CD29 is present on a large proportion of EVs, alone, or together with CD9 and/or with CD63, while the 3 other markers (CD44, SSEA-4, CD98) are not as frequently detected on EVs, are often together with CD9 and mainly on EVs of larger size.

### Modulation of the composition of MDA-MB-231-EVs by drug treatment of the corresponding cells

Finally, we asked whether our pipeline of single EV analysis could help the search for tools that modulate specifically the release of EV subtypes. We have previously screened a large library of (mostly) FDA-approved drugs, for their effect on EV release (Grisard *et al*., 2022), using nanoluciferase enzyme-tagged CD63 or CD9 to quantify secretion of EVs bearing one or the other tetraspanin directly in the CM of cells. Validation of the screen results could only be done at that time by counting particles, or bulk analyses of proteins of the EVs released by the drug-treated MDA-MB-231 cells: western blot or multiplexed bead-based EV capture. We selected three drugs which either increased (Homosalate) or decreased (Dipivefrin, Metaraminol) the release of nanoluciferase-tagged CD9 and/or CD63 on EVs and one drug (Ebselen) which displayed inconsistent effects on EV release measured by nanoluciferase activity (decrease in (Grisard *et al*., 2022)) versus bulk EV quantification (increase: Fig. 4A, right panel). MDA-MB-231 cells were treated for 24 hours with DMSO (solvent control) or each drug, after which the CM containing EVs was collected, and cell viability was determined. EVs were isolated by SEC and analyzed by Flow NanoAnalyzer for particle count, size and expression of CD63 and CD9. Of note, the drugs did not impair cell viability, which remained between 94% and 98.5% (Fig. 4A, left panel). We observed that the EV size was slightly reduced with Dipivefrin (68.1±1.1 nm) and Metaraminol (68.3±1.7 nm) and slightly increased with Ebselen (73.1±0.9 nm) compared to the DMSO control (70.8±1 nm) (Fig. 4A, middle panel). As compared to the DMSO control, Homosalate and Ebselen increased 2.26±0.49 and 1.96±0.33 times, respectively, the number of EVs released by a given number of plated cells, while Dipivefrin and Metaraminol led to production of fewer EVs (0.76±0.05 and 0.88±0.03 times, respectively) (Fig. 4A, right panel). To verify that these modifications in EV release were not due to changes in cell number or adhesion induced by the drugs, we also measured cell behaviour, both in the absence (DMEM or DMSO) or in the presence of the drugs (diluted in DMSO), using the xCELLigence system (Sup. Fig. 5A). This system measures continuously adhesion and number of cells, and calculates a “cell index” as a proxy for cell growth. Under the same culture conditions used for EV isolation (16 hours with DMEM + 10% FCS followed by 24 hours with DMEM +/- drugs), we observed that cells treated with Dipivefrin and Metaraminol exhibited very slightly increased cell index, cells treated with Ebselen showed slightly decreased index, and no change was observed for Homosalate treatment, compared to DMSO condition. Therefore, the observed effect of these drugs on cell index was negligible and, when existing, inverse to their effect on EV release, ruling out that the drugs changed EV number because they changed cell growth. To know if these changes in total EV number also resulted in changes in proportions of EV subtypes, we analyzed the expression of CD63 and CD9 (Fig. 4B, Sup. Fig. 5B). Interestingly, Homosalate, compared to the DMSO condition, proportionally decreased the fraction of CD63+CD9-EVs and increased the fraction of CD63+CD9+ EVs (Fig. 4B, and raw data in Sup. Fig. 5B). In comparison, although Ebselen increased EV release similarly to Homosalate, it did not alter the distribution of CD63 and CD9 on the surface of the EVs. Finally, a different pattern of alteration was observed with Dipivefrin and Metaraminol, which increased the fraction of CD63+CD9-EVs and decreased the fraction of CD63-CD9+ EVs, without affecting the proportion of CD63+CD9+ EVs. Overall, single EV analysis by the Flow NanoAnalyzer clarifies the effects of drugs on the secretion of EV subtypes from treated cells.

**Figure 4:**
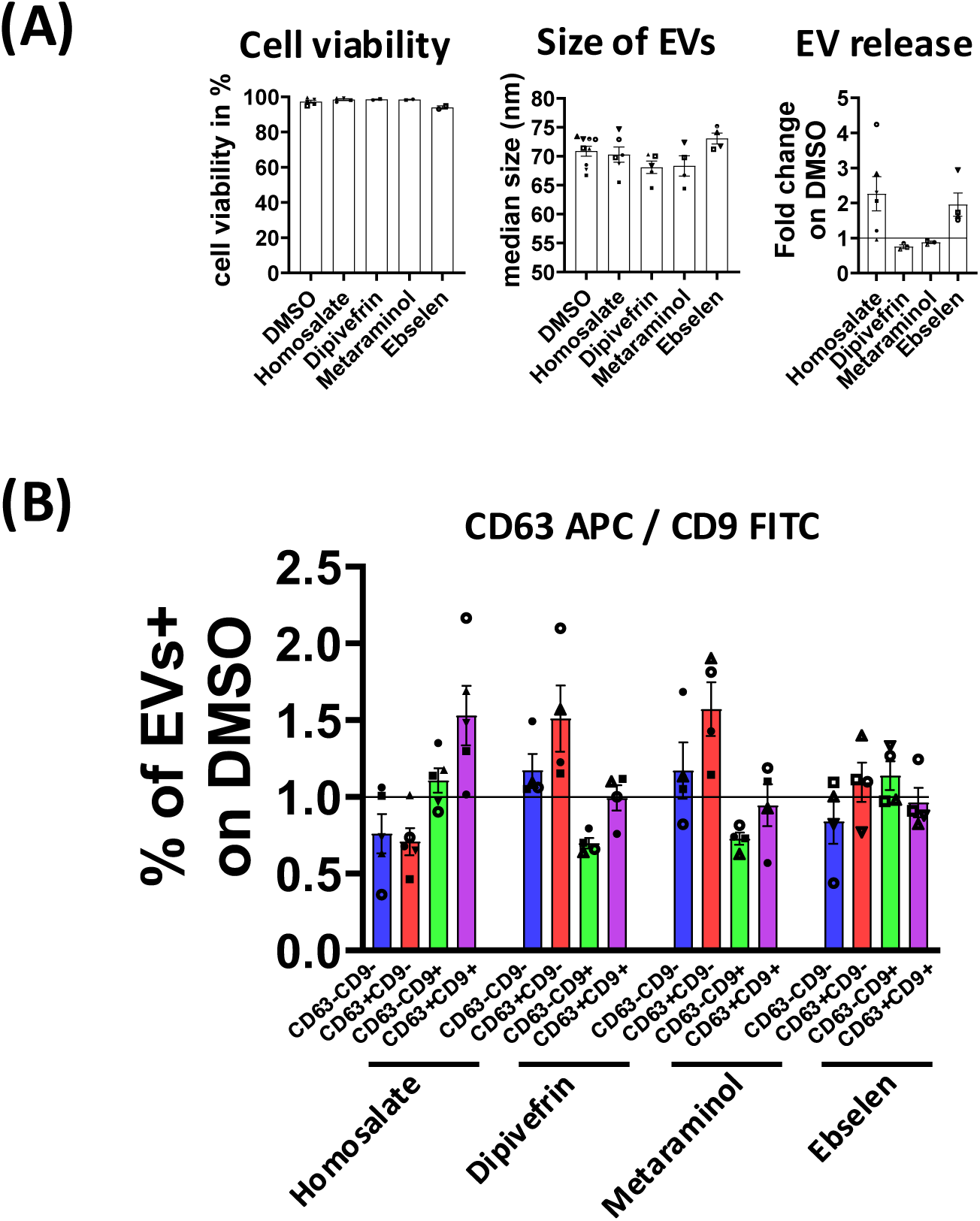
Different drugs affect differently the proportion of subtypes of EVs released by MDA-MB-231. A) Quantification of cell viability (left panel) after Homosalate, Dipivefrin, Metaraminol or Ebselen (10 µM each) treatment of MDA-MB-231 cells, as compared to control (DMSO) treated cells (mean ± SEM of n=2-5 independent experiments). Median size (middle panel) of MDA-MB-231-EVs isolated after SEC and measured by Flow NanoAnalyzer (mean ± SEM of n=4-9 independent experiments). Quantification of EV release after MDA-MB-231 treatment compared to control (DMSO) (right panel). The total number of EVs (quantified by Flow NanoAnalysis) released by treated cells was divided by the total number of EVs released by control cells (DMSO) for the same number of plated cells (mean ± SEM of n=3-6 independent experiments). B) Double staining CD63-APC/CD9-FITC of MDA-MB-231-EVs after drug treatment. Bar charts showing the percentages of double-negative (blue), single-positive (red: APC, green: FITC) and double-positive (purple) EV populations released by drug-treated MDA-MB-231 cells compared to control cells (DMSO) in the same experiment (mean ± SEM of n=4-5 independent experiments). Raw data are shown in Sup. Fig. 5B.

## DISCUSSION

The work presented here describes 1) a specific and sensitive protocol to label EVs with fluorescent antibodies before analysis by a nanoparticle-optimized flow analyzer, 2) its use to uncover the variability in proportions of EV subtypes released by different cell lines, and 3) its use to uncover the changes in subtypes of EVs released by a given cell line exposed to different drug treatments. Our work thus paves the way for exhaustive evaluation of the heterogeneity of EVs recovered from any source, and its modification induced by change in the environmental condition of the producing cells.

Although other groups had previously used a previous or this version of the nano flow cytometer to quantify the EV-enriched tetraspanins (CD9, CD63, CD81) on tumor-derived EVs (Brealey *et al*., 2024; Lees *et al*, 2022; Tian *et al*, 2018), the published protocol for EV staining was not detailed enough to allow direct implementation. We thus developed here a protocol that includes two major features to reach a high level of specificity and sensitivity: respectively, the use of antigen-negative EVs as negative controls, and the elimination of excess or non-specifically attached fluorescent antibodies by size-exclusion chromatography (SEC). The former is crucial to determine the threshold of positivity and was instrumental in showing that simple dilution of the excess antibody was not sufficient to allow specific and sensitive identification of antibody-labeled EVs (Fig. 1). Indeed, upon dilution of the antibody, the best difference of staining observed between WT and CD81 KO EVs was about 20%, whereas it raised to about 40% when all background was eliminated. Published articles from the instrument provider showing results obtained upon post-staining sample dilution (Brealey *et al*., 2024; Lees *et al*., 2022) show indeed differences in signals depending on the antibody dilution, comparing EVs with a “background” control. However, the nature of this control was not clearly described, and it was probably based on the background signal observed in buffer with antibodies, thus not taking into account the potential unspecific binding of the antibody to EVs. We show here that the optimal antibody concentration can only be properly identified comparing antigen-expressing EVs to negative control EVs not bearing the epitope recognized by the antibody. As we did here for the anti-tetraspanin stainings, using EVs from cells KO for the molecule of interest is the best option, but it is not always achievable for all new antibody targets. In that case, we propose to use EVs from another species, of which the antibody does not cross-recognize the antigen, as negative controls. This is not the optimal situation, since EVs from another cell source could theoretically display different intrinsic background staining, but it is the best option we could come up with. Mouse cell-derived EVs were appropriate for the 4 other antibodies we also used here (ITGB1/CD29, CD44, SSEA-4, SLC3A2/CD98), but in other cases, the anti-human antigen-antibody may recognize the mouse epitope. In this condition, other EVs from more distant species (e.g. chicken, hamster) or from cells known to not express the antigen (if they exist) will have to be produced.

Such negative control EVs are also instrumental in identifying methods to eliminate free antibodies after staining. Here we used SEC-based columns, as described before by others (Morales-Kastresana *et al*, 2017), and successfully used by us to eliminate free amine-binding fluorescent dye post-EV staining (Loconte *et al*, 2023), but with a miniaturized commercial version, allowing to save time in the process. We noticed the loss of about 50% of particles upon mini-SEC cleaning, but all particles of 30-250 nm size, with or not antibodies bound to their surface, should interact similarly with the neutral beads packing the column. Indeed, we did not observe any difference in size of HeLa EVs recovered after mini-SEC cleaning, thus there is no bias in the final analyzed EV population introduced by this step, at least for the EVs analyzed here. Faster or easier methods, like filtration on 10 kDa pore filters, which worked for the elimination of a lipid dye post-EV staining (Loconte *et al*., 2023), or ultracentrifugation of EVs post-labeling did not work in our hands to eliminate unbound antibodies and/or induced EV aggregation, but we cannot exclude that other methods may be eventually suitable.

In conclusion, each new antigen to be analyzed by Flow NanoAnalysis will require initial determination of the proper negative control EVs, optimal antibody dose and cleaning process. This would apply not only to the instrument used here, but also to all other nanoparticle-dedicated flow cytometers. For the antibody titration step, it should ideally be performed on each new source of analyzed EVs, like titration is done in regular flow cytometry on each analyzed cell, to account for potential differences in intrinsic background fluorescence of different cells. However, we expect EVs to display limited intrinsic fluorescence and, at least for non immune cell-derived EVs, to not bear generic antibody-binding molecules (such as Fc receptors), thus to not display different background antibody signals. Indeed, we obtained the same optimal dose of anti-tetraspanin antibodies on 2 different EVs versus their respective controls (HeLa and MDA-MB-231, Sup. Fig. 2), and thus we used this dose on the third source of EVs (A549), without re-performing a titration on these EVs.

As a characterization method of heterogenous EVs, the nanoparticle-optimized Flow NanoAnalyzer used here (similarly to other nanoparticle-dedicated flow analyzers) has the strong advantage over other fluorescence methods that it detects all particles (larger than about 40 nm in diameter) by their side-scatter properties with the 488 nm laser, without the need for another fluorescent label. It thus does not exclude any EVs, which other methods do. For instance, another fluorescence- and flow-based instrument is the imaging flow cytometer (ImageStreamX, Amnis). However, before any analysis, this instrument requires EVs to be labeled with a fluorescent tag, since EVs are defined as events non-detectable in the bright-field channel, but detectable in one of the fluorescence channels. So far, the strategy to fluorescently label EVs is achieved either by various dyes (with the caveats of false positive events due to the dye itself and of unknown efficiency of label) (Lannigan & Erdbruegger, 2017), or by genetically engineering EV-producing cells to make their EVs express CD63-eGFP (Gorgens *et al*, 2019), or by staining EVs with antibodies against tetraspanins (Ricklefs *et al*, 2019) or other surface molecules (Holcar *et al*, 2025). In all cases, therefore, a variable proportion of non-labeled EVs is ignored by this method and the calculated proportions of single- or double-positive EVs do not apply to the total EV population. Therefore, further developments of EV analysis by imaging flow cytometry require identification of a universal method for fluorescence staining of EVs, which may ultimately be achieved by comparison with other single EV analysis methods such as the Flow NanoAnalysis (Brealey *et al*., 2024).

On the other hand, other approaches use regular immunofluorescence (combined with interferometric reflectance (Daaboul *et al*, 2016), or with limited diffraction (Schurz *et al*, 2022)) or super-resolution microscopy (Saftics *et al*, 2023) to analyze single EVs. However, this requires pre-capture of EVs on a slide, which is achieved in a vast majority of approaches by capturing EVs via antibodies recognizing CD9, CD63 and/or CD81, separately or together. Since our analysis shows in 3 different cell lines that at least 20% (HeLa, MDA-MB-231) and up to 40% (A549) of their EVs do not bear any of these 3 tetraspanins to a level allowing their detection by Flow NanoAnalysis, these capture-based immunofluorescence instruments analyse only a subpopulation of EVs. Our observations on 3 human breast, cervical and lung carcinoma cells are consistent with those of other groups using also Flow NanoAnalysis, on 2 colon carcinoma cell lines (60% or 40% of EVs negative for the 3 tetraspanins in (Lees *et al*., 2022)), although these results were obtained without antibody elimination, thus reducing the specific signal and possibly increasing the % of negative EVs. Another study using successive cycles of EV antibody staining/destaining in a microfluidic chip also concluded that between 25 and 53% of EVs did not bear CD9/CD63/CD81 in two pancreatic cancer cell lines (Spitzberg *et al*, 2023). Therefore capture via anti-tetraspanin antibodies introduces a bias, and no pan-EV capture tools have yet been confidently identified, but they will be necessary to provide a non-biased picture of the analyzed EVs. A difficulty, however, will be to demonstrate that all EVs, whatever their size or their surface determinants, are equally captured.

Finally, our experimental results using the improved Flow NanoAnalysis protocol show the power of single EV analysis to demonstrate hypotheses made from observations of semi-quantitative bulk (e.g. Western blot) or semi-bulk (e.g. MacsPlexEV) previous analyses of EVs. First, we show that, even when originating from a single *in vitro* cultured cell line, i.e. the most homogenous possible source, EVs are very heterogeneous in terms of exposed surface molecules. In addition to the CD9, CD63 and CD81 tetraspanins, we identified CD29 (the integrin beta 1 chain, ITGB1) as another very widely present surface marker of EVs, associated or not with the tetraspanins, worth exploring further for its presence on many other sources of EVs. Conversely, CD81 was unexpectedly observed only on a small minority of MDA-MB-231 EVs (4% or less), even though previous Western blot and MACSPlexEV analyses detected CD81 apparently similarly to CD63 or CD9 (Grisard *et al*., 2022). For the MACSPlexEV analysis, it semi-quantifies EVs bearing the molecule recognized by the capture beads together with either one of the 3 tetraspanins, or with CD98 (Sup. Fig. 4): the actual measured Median Fluorescence Intensity (MFI) depends on the affinity and fluorescence of each antibody and thus provides a hint on the level of expression, but not an actual demonstration. The Flow NanoAnalysis does not suggest the total absence of CD81 on MDA-MB-231, but rather the presence of a very minor subtype, primarily together with CD9 (Fig. 3D). This could be due to a very low number of CD81 molecule on each EV, many of them falling below the detection threshold of the Flow NanoAnalyzer, or alternatively, to a particular conformation, such as binding to different protein partners or altered glycosylation, not observed on EVs from HeLa nor A549, making the epitope inaccessible to antibodies. Masking of the epitope recognized by the clone 5A6 has indeed been observed when CD81 is associated to CD19 in B cells (Susa *et al*, 2020). Another protein partner of CD81 in MDA-MB-231 EVs could thus have a similar effect here.

Finally, our results on the distribution of each tetraspanin on subpopulation of EVs are very informative. When comparing the 3 cell lines, we noticed that CD9 was the most prevalent tetraspanin detected on HeLa and A549 EVs, on almost 3 times more EVs than CD63, while equal proportions of EVs bore CD9 and CD63 in MDA-MB-231 (Fig. 2A). Furthermore, a higher proportion of EVs from MDA-MB-231 (around 30%), than from HeLa or A549 EVs (respectively around 20% and 10%) bore CD9 and CD63 together. This could be correlated with the subcellular distribution of these molecules: as we (Mathieu *et al*., 2021) and others (Fordjour *et al*, 2022; Rous *et al*, 2002) described before, in most cells at steady state CD63 accumulates in late endosomes, this is why it is generally considered as an exosome marker, while CD9 accumulates at the plasma membrane, and thus qualifies as marker of ectosome. However, by performing staining for the 3 tetraspanins in cells (Sup. Fig. 7), we observed that, even if this global distribution was indeed observed in the 3 cells used here, MDA-MB-231 clearly displayed some CD63 colocalized with CD9 at the plasma membrane. The size of ectosomes can be anywhere between the size of exosomes (which matches the size of intraluminal vesicles of MVBs) and much larger, therefore, since we observed CD63+ EVs primarily larger than the median size of MDA-MB-231 EVs whereas in HeLa cells CD63+ EVs were mainly smaller than median size (Fig. 2B), our results suggest that MDA-MB-231 probably release more ectosomes bearing CD63 than the other cell lines, in addition to bona fide exosomes. The other 3 surface molecules, CD44, SSEA-4 and CD98, mainly released on large EVs (Fig. 3B), are therefore also likely associated with ectosomes. Furthermore, the change of tetraspanin profile observed on EVs released by Homosalate-treated cells (Fig. 4), with more CD63+/CD9+ but less CD63+/CD9-EVs, is likely due to an increased release of multi-tetraspanin-bearing ectosomes, while, conversely, Dipivefrin and Metaraminol increase the proportion of CD63+/CD9-EVs, thus favour release of *bona fide* exosomes, to the expense of ectosomes. Flow NanoAnalysis also clearly demonstrate that Ebselen is another drug promoting EV release, but, different from Homosalate, it induces the release of all types of EVs from any subcellular origin, both ectosomes and exosomes (Fig. 4). Finally, CD81 is distributed in the 3 cell lines both at the plasma membrane, and in intracellular compartments distinct from those containing CD63 (Sup. Fig. 7). In HeLa and MDA-MB-231, CD81 is exclusively detected on EVs bearing also CD9 (Fig. 2D), suggesting that it is secreted on ectosomes. In MDA-MB-231, CD81 accumulates more in large internal compartments than at the plasma membrane, which could also contribute to explain why it is detected on very few EVs, if these large compartments were not competent for secretion of their content. By contrast, A549 cells release a significant portion of EVs bearing CD81 without CD9 nor CD63: it would be worth exploring whether these CD81+ EVs come from the CD63-negative intracellular compartments, thus representing another type of “exosomes”, maybe corresponding to those induced by stress in other cells (Fan *et al*, 2020).

Overall, the observations obtained thanks to the unbiased, sensitive, and specific single EV analysis method described here open the door to extremely powerful basic science cell biology studies of the mechanisms of EV biogenesis, but also to improvements of clinical applications, by allowing identification and ultimately selection of the few EVs bearing relevant therapeutic compounds among the bulk population of EVs released by the producing cells. However, as with any novel technology, some so far un-identified biases or artefacts may eventually be found in Flow NanoAnalysis, which will have to be carefully solved, as we did here, by using relevant negative controls.

## MATERIALS AND METHODS

See Sup. Table 2 (excel file) for a reporting check-list according to MIFlowCyt-EV (Welsh *et al*., 2020) and MIFlowCyt, following the model downloaded from https://www.evflowcytometry.org/resources/.

### Cell culture

HeLa and MDA-MB-231 cells were cultured in DMEM-Glutamax (Gibco), and A549 in RPMI-1640-Glutamax (Gibco), all with 10% Fetal Calf Serum (FCS, Gibco), 100 U/ml penicillin and 100 μg/ml streptomycin (Gibco). The mouse cells E0771 were cultured in RPMI-1640-Glutamax, with 10% FCS (Gibco), 20 mM HEPES, 10 mM sodium pyruvate, 100 U/ml penicillin and 100 μg/ml streptomycin (Gibco). Cell lines were grown at 37°C, under 5% CO2, in humidified incubators and routinely tested using Mycoplasma detection Kit (Eurofins) for mycoplasma contamination. Only mycoplasma negative cells were used for experiments. Human cells were validated by short-tandem repeat (STR) in 2018 (HeLa, MDA-MB-231) and 2025 (MDA-MB-231, A549). HeLa cells KO for CD9 or CD63 generated by CRISPR/Cas9 method, and used as a pool of 5 clones obtained after two rounds of sorting (CD9, (Mathieu *et al*., 2021)) or as one sub-clone from the bulk population after 2 rounds of sorting (CD63 (Palmulli *et al*., 2024)) have been previously described. HeLa cells KO for CD81 were generated by CRISPR/Cas9 (guide RNA sequence: AGGAATCCCAGTGCCTGCTG) and used as a pool of 38 subclones.

### EV isolation and drug treatment

Cells were seeded in an optimized number to obtain a confluency of 80-90% on the day of EV isolation (6 × 10^6^ cells per 15 cm culture dish). After 16 h, cells were washed 1X with PBS and 25 ml of serum-free medium were added per 15 cm culture dish. For drug treatments, we added to serum-free medium 25 μl of DMSO (0.1% final), or 25 μl of stock solutions at 10 mM in DMSO of Homosalate, Dipivefrin hydrochloride, Metaraminol bitartrate or Ebselen (GreenPharma Labs), leading to 10 μM final. Twenty-four hours later, CM was harvested and centrifuged at 400 g for 10 min at 4°C to remove dead cells and debris. In parallel, cells were detached and the percentage of cell viability was determined using erythrosin B (Sigma). The CM was then centrifuged at 2,000 g for 20 min at 4°C to discard large EVs in the 2K pellet and then concentrated on a Centricon Plus-70 Centrifugal Filter (MWCO =10 kDa; Millipore, #UFC701008). CM concentrated to 500 μl was loaded on 35 nm qEV size-exclusion columns Gen 2 (Izon) for separation. After discarding the void volume and according to the manufacturer’s instructions, each SEC fraction was collected in 400 μl volume. For experiments, we collected SEC fractions 7 to 11 (F7-F11) corresponding to EV-rich fractions and pooled fractions were further concentrated using 10 kDa cut-off filters (Amicon Ultra-15, Millipore). Then, concentrated SEC fractions were individually analyzed in terms of particle number and particle size using nano-flow cytometry (U30 Flow NanoAnalyzer, NanoFCM) before staining. This protocol of EV isolation is performed routinely in our laboratory, and we have extensively characterized the resulting MDA-MB-231 EVs by western blotting and transmission electron-microscopy in our previous work (Grisard *et al*., 2022; Loconte *et al*., 2023; Tkach *et al*, 2022), for which EV-TRACK (Roux *et al*, 2020) record had been filed.

### Staining of EVs with antibodies for nano-flow cytometry

Concentrated EVs obtained after SEC were diluted at 2-3.10^10^ particles/ml in PBS and then 1 µl of a given dilution of antibody in PBS, for the single staining, or 1 µl of each antibody for the double staining, was added to 9 μl of EVs in suspension. Antibodies and EVs were incubated for 30 min at 4°C. For the initial condition of “stained EVs diluted in PBS” (Fig. 1A), the samples were diluted 100 times with PBS after the incubation and EVs were directly analyzed on the U30 Flow NanoAnalyzer. For the following conditions of “stained EVs washed on SEC column” (Fig. 1B and all subsequent experiments), each 10 µl mix of EVs and antibodies was diluted 10 times with PBS and applied to the top of an Exo-spin Mini SEC column containing beads with around 30 nm pores (Interchim). The void volume was discarded and the EVs were recovered in 180 µl of PBS and analyzed directly by Flow NanoAnalyzer.

Before analyzing labeled vesicles, we first checked if the antibodies formed aggregates detected by the Flow NanoAnalyzer. For that, 1 µl of antibody (at a final concentration of 4 or 20 µg/ml for anti-CD63 and anti-CD81, 2 or 10 µg/ml for anti-CD9) was added to 9 µl of PBS and incubated for 30 min at 4°C. The samples were diluted 100 times with PBS after the incubation and analyzed with the U30 Flow Nanoanalyzer. We observed that anti-CD63 APC, anti-CD81 APC, anti-CD81 FITC and anti-CD9 FITC antibodies formed few rare fluorescent aggregates between 55 and 110 nm, which were eliminated by mini-SEC cleaning (Sup. Fig. 6A). In the experimental condition at 20 µg/ml which corresponds to the highest antibody concentration, we observed no interference of free antibodies which could affect the analysis of labeled EVs detected by Flow NanoAnalyzer. For control purposes, non-stained HEK293-FT EVs and CD81 APC- or CD81 FITC-labeled EVs analyzed with the Flow NanoAnalyzer were then lysed by NP40 (1%) for a minimum 30 min at RT and re-analyzed with the same settings (HEK293-FT were cultured in DMEM-Glutamax (Gibco) with 10% Fetal Calf Serum (FCS, Gibco), 100 U/ml penicillin and 100 μg/ml streptomycin (Gibco) and EVs isolated after 24h of production in medium with EV-depleted serum as we did previously (Cocozza *et al*, 2020b)): 90% of CD81-APC- and 60% of CD81-FITC-stained EVs disappeared after detergent treatment, and we also observed that NP40 at 1% formed micelles detected as events by the Flow NanoAnalyzer (Sup. Fig. 6B). Finally, serial dilution of CD63-APC- and CD9-APC-labeled MDA-MB-231 EVs were done to discard swarm effect and show that single EVs are actually analyzed at the concentrations used for the analysis with no change of the Median Fluorescence Intensity (MFI) observed in the different dilutions (Sup. Fig. 6C).

### Nano-flow cytometry of EVs

EVs were analyzed using a U30 Flow NanoAnalyzer (NanoFCM) equipped with two lasers (488 nm and 638 nm) and five band-pass filters (488/10, 525/40, 580/40, 670/30 and 710/40). All measurements were performed according to the manufacturer’s protocol. Briefly, a silica nanoparticle cocktail (68–155 nm, Cat. # S16M-Exo, NanoFCM) was used as the size reference and polystyrene 250 nm beads (QC Beads, Cat. # S08210, NanoFCM) for particle concentration, light scatter and fluorescence calibration, and to validate the flow rate of the machine. The PBS sample buffer was run to set the background threshold. Diluted EVs in PBS were run at a flow rate of ± 30 nl/min leading to 50-170 events per second, with between 3,000 and 10,000 events recorded over 60 seconds. All samples were run under identical pressure (1 kPa), and the signals of SSC, FITC and APC were recorded. Height (H) values of these signals were used for analyses. Antibodies used for the staining are listed in Sup. Table 1. The analysis and graphs were obtained with the FlowJo v10 software. Gate for negative events was defined on unstained EVs, and applied to stained EVs, where the Median Fluorescence Intensity (MFI) of positive and negative events was calculated. For each anti-tetraspanin antibody (CD63, CD81, CD9 APC or FITC), a serial dilution with four different concentrations was first tested on the HeLa WT and KO EVs. The stain index determined for each antibody and each dilution was calculated as follows on the HeLa WT EVs: (MFI of Positive population - MFI of Negative population) / (SD of Negative population X 2) and is shown in Sup. Fig. 2. The optimal antibody concentration was chosen based on the highest stain index while providing a negative staining for the HeLa KO EVs. Then, the same antibodies were tested on the MDA-MB-231 and E0771 EVs. For this, the optimal antibody concentration was chosen based on the highest stain index for MDA-MB-231 EVs while providing a negative staining for the E0771 EVs. We observed that the optimal antibody concentrations chosen for HeLa and MDA-MB-231 EVs were identical. For the anti-CD29 (AF488 or AF647), anti-CD44 (AF647), anti-SSEA-4 (APC) and anti-CD98 (APC) antibodies, a serial dilution with four or five different concentrations was tested only on MDA-MB-231 and E0771 EVs. In the same manner, the optimal concentration chosen for each antibody was the one that provided the highest stain index for MDA-MB-231 EVs while giving negative staining for E0771 EVs. Size parameter (y) was created in FlowJo using the formula y = ax^(b)+c with the a, b, c parameters obtained after calibration of size (x) with silica 68-155 nm beads. Raw data as .fcs files will be submitted to http://flowrepository.org/ before final publication of the article.

### Imaging flow cytometry of cells

1.10^6^ Hela WT or KO cells were labeled with fixable viability dye efluor 780 (eBioscience) for CD9 SB436 labeling or BD Horizon™ Fixable Viability Stain 450 (FVS450) for CD63 AF594 or CD81 AF647 labeling at a dilution of 1/1000 per volume. After the labeling, we performed three successive steps of washing by centrifugation at 300 g for 10 min with flow cytometry buffer (0.5% BSA, 2mM EDTA in PBS). A supplementary step of fixation/permeabilization using the FIX & PERM™ Cell Permeabilization Kit (Invitrogen) was done for CD63 labeling following the manufacturer’s instructions in order to detect intracellular CD63 staining. Antibody staining was done in a volume of 100 µl with anti-CD9-SB436 (eBioscience, # 62-0098-42, clone SN4) at a concentration of 0.25 µg/million cells, and with anti-CD81-AF647 (Santa Cruz Biotechnology, # sc-166029, clone 5A6) and anti-CD63-AF594 (Biolegend, # 353034, clone H5C6) at a concentration of 1 µg/million cells. Isotype controls with the same fluorochromes were used to determine the positive events: SuperBright 436 (eBioscience, # 62-4714-82), Alexa Fluor 647 (Santa Cruz, # sc-24636) and Alexa Fluor 594 control (Biolegend, # 400174). Samples were analyzed by imaging flow cytometry (ImageStream X MKII, Amnis/Luminex) with the following settings: objective 60X, low speed of fluidics, brightfield and side scatter ON. The automatic compensation wizard was used, to create a compensation matrix on single-labeled sample. Analysis was done using IDEAS software (v6.2), see illustration of gating in Sup. Fig. 1A. We first selected ‘Singlet’ cells (Ch01 Area and Aspect Ratio Intensity), then dead cells and non-circular cells (‘Live Circular’) were excluded using respectively dye staining (intensity LD450) and Circularity feature on brightfield, finally focused events were selected in ‘Focus’ (Ch01 Gradient RMS).

### Nanoparticle-tracking analysis of EVs

For EV analysis by MACSPlexEV, EVs were quantified by nanoparticle tracking analysis (NTA) instead of Flow NanoAnalyzer, to match the conditions of our previous work (Grisard *et al*., 2022). NTA was performed using ZetaView PMX-120 (Particle Metrix) equipped with a 488 nm laser, at 10X magnification, with software version 8.05.02. The instrument settings were 22°C, gain of 26 and shutter of 70. Measurements were done at 11 different positions (two cycles per position) and frame rate of 30 frames per second. Image evaluation was done on particles with Minimum Brightness: 20, Minimum Area: 10, Maximum Area: 500, Maximum Brightness: 255. Tracking Radius2 was 100, Minimum Tracelength: 7.

### EV analysis by bead-based multiplex flow cytometry assay

Concentrated EVs were processed according to manufacturer instructions to perform bead-based multiplex analysis by flow cytometry (MACSPlexEV Kit, human, Miltenyi): 1.10^9^ MDA-MB-231-EVs (quantified by nanoparticle-tracking analysis, corresponding to approximately 3-5.10^8^ EVs quantified by Flow NanoANalyzer) were diluted with MACSPlex buffer to a final volume of 120 μl and 15 μl of MACSPlex Exosome Capture Beads were added. Samples were incubated on an orbital shaker overnight at room temperature protected from light. After washing, detection antibodies (APC-conjugated anti-CD9/anti-CD63/anti-CD81 mix at dilution recommended by the provider, or 5 μl anti-SLC3A2/CD98-APC (clone REA387, Miltenyi) were incubated for 1 h at RT, before washing twice in 500 μl buffer. Flow cytometric analysis was performed with Aurora Analyzer (Cytek) and data analyzed with FlowJo software (v10, FlowJo LLC). The 39 single-bead populations were gated to allow the determination of APC signal intensity on the respective bead population. Median fluorescence intensity (MFI) for each capture bead was divided by the respective MFI values from matched non-EV controls that were treated exactly like EV-containing samples (including incubation with detection antibodies).

### Cell growth analysis by live impedance measurement

MDA-MB-231 cells were resuspended in DMEM with 10% FCS (150 µl/well), transferred into xCELLigence microplates (E-plate 16, Agilent) at a density of 20,000 cells per well and plates were loaded into xCELLigence RTCA DP instrument (Agilent) inside a 37°C incubator for 16h. The run was then paused, cells were washed 1X with PBS and then incubated with DMEM without FCS (condition « no treatment »), DMSO or drugs. A run of 24 h with readings of impedance every 30 min was programmed. Cell-sensor impedance is expressed as an arbitrary unit called the Cell Index (CI).

### Immunofluorescence analysis of cells

One day before staining, 100,000 HeLa, MDA-MB-231 or A549 cells were seeded on 12 mm diameter coverslips coated with poly-l-lysine (15 μg/ml, Sigma). On the day of staining, cells were fixed with 4% paraformaldehyde (PFA) (EMS) for 15 min at RT and then washed 3 times with PBS 1X. The primary and secondary antibodies were successively incubated for 1 h each at RT in PBS containing 0.1% saponin and 0.1% BSA. Primary antibodies were: mouse IgG2b anti-human CD63 (clone TS63b, 1/100, available upon request to E. Rubinstein), mouse IgG2a anti-human CD81 (clone TS81, 1/100) and mouse IgG1 anti-human CD9 (clone TS9, 1/100) (commercially available at Diaclone or Abcam). Secondary antibodies were: goat anti-mouse IgG2b Alexafluor 647 (Invitrogen, 1/200), goat anti-mouse IgG2a Alexafluor 568 (Invitrogen, 1/200) and goat anti-mouse IgG1 Alexafluor 488 (Invitrogen, 1/200). Coverslips were then mounted on slides with Fluoromount G (Invitrogen). Images were acquired on a Zeiss LSM 780 confocal microscope using an alpha Plan-Apochromat 63x/1.46 Oil. On each slide, laser powers were set to avoid saturation.

## ACKNOWLEDGEMENTS

We thank Drs R. Palmulli and G. van Niel (INSERM U1307/CNRS UMR6075, Nantes, France) for providing the clone of CD63 KO HeLa cells. We thank Louise Merle and Ahmad Maswadeh for helpful comments on the manuscript.

This work was funded by INSERM, CNRS, Institut Curie, SIRIC Curie/INCa-DGOS-Inserm-ITMO Cancer_18000, PSL research University, grants from french ANR (ANR-22-CE18-0012), INCa (INCa_16083, INCa_16735), Fondation ARC (PGA12021020003189_3588), Fondation Chercher et Trouver, and ITMO Cancer of Aviesan within the framework of the 2021-2030 Cancer Control Strategy, on funds administered by Inserm for purchase of the U30 Flow NanoAnalyzer.

We also acknowledge the following Core facilities of Institut Curie: Cell and Tissue Imaging (PICT-IBiSA), member of the French national research infrastructure France-BioImaging (ANR10-INBS-04) for fluorescence microscopy, Genomic for STR-based validation of human cells, Cytometry for assistance in data acquisition and imaging flow cytometry analysis.

---

**Sup. Figure 1:**
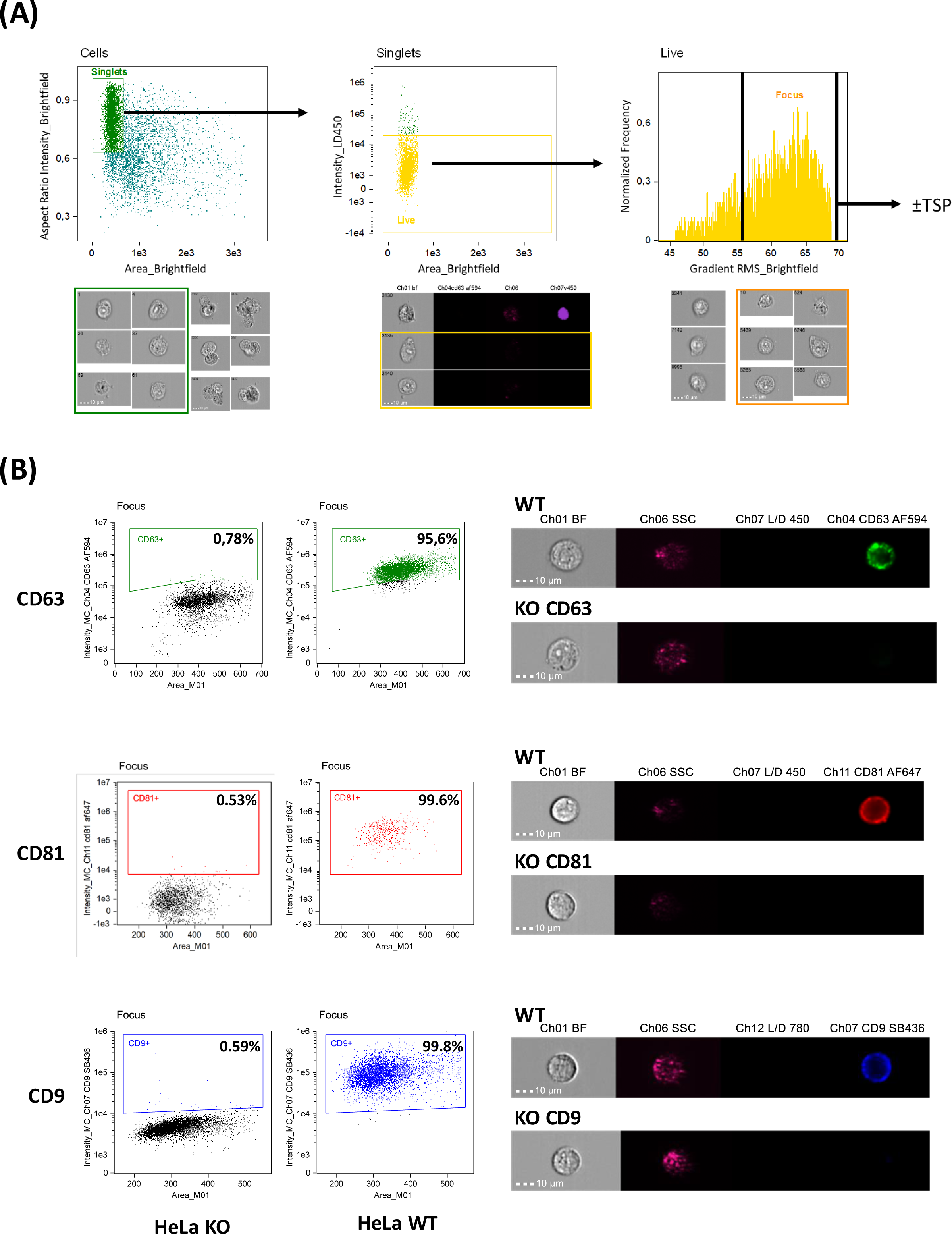
Imaging flow cytometry validation of Hela CD63, or CD81, or CD9 KO cells compared to Hela WT cells. A) Gating strategy for ImageStream analysis. Single cells were first selected as ‘Singlet’ cells (Ch01 Area and Aspect Ratio Intensity), then dead cells and non-circular cells (‘Live Circular’) were excluded using respectively dye staining (intensity LD450) and Circularity feature on brightfield, finally focused events were selected in ‘Focus’ (Ch01 Gradient RMS). The percentage of positive cells for the different tetraspanins (TSP) was determined in focused lived single cells. Representative dot plots of the gating strategy (upper panel) and representative images of the gated cells (lower panel) are shown. B) 1.10^6^ Hela WT or KO cells for each of the tetraspanins were labeled with fixable viability dye and washed before antibody staining. A supplementary step of fixation/permeabilization was done for CD63 labeling. The gate for positive events for CD9-SB436, CD81-AF647 or CD63-AF594 was established with isotype controls with the same fluorochromes. Percentage of tetraspanin+ cells for Hela KO for CD63, CD81 or CD9 compared to Hela WT cells were determined. Illustration of dot plots of Area/Intensity (left) of WT cells labeled with the isotype control (black) or the specific antibody (colour), and images of individual HeLa WT or KO cells (right).

**Sup. Figure 2:**
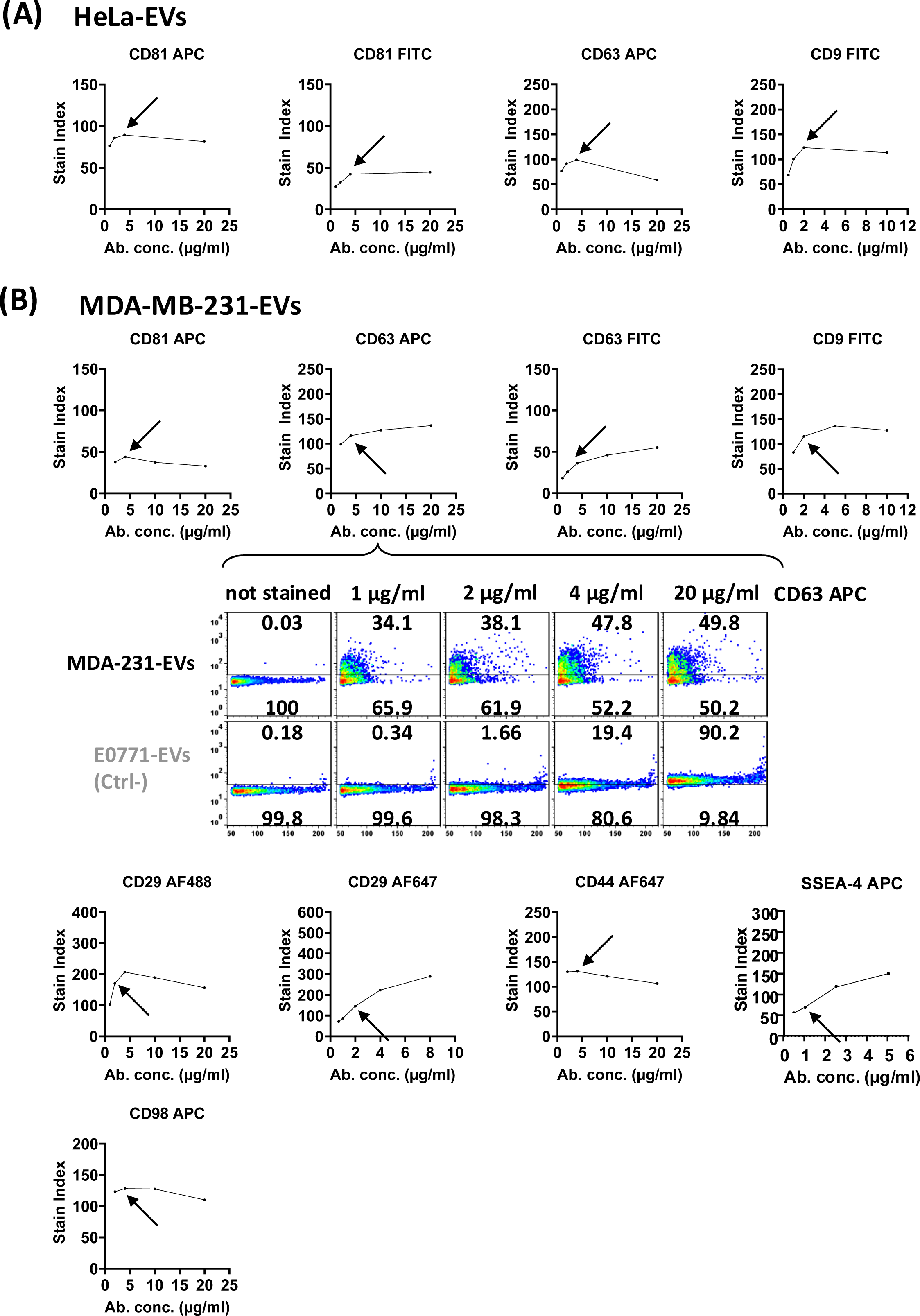
Antibody titration on EVs. A-B) For each antibody and batch, plotting the stain index for each concentration of antibody allowed to titrate the optimal amount of antibody used with EV staining. For each antibody, 4 or 5 different dilutions were performed, and the stain index for each dilution on the antigen-expressing EVs was calculated according to the following formula: Stain Index = (Median Fluorescence Intensity (MFI) of Positive population - MFI of Negative population) / (SD of Negative population x 2). Gate for negative events was defined on unstained EVs, and applied to stained EVs, where MFI of negative and positive EVs was separately determined. The lowest concentration reaching the maximum level of stain index combined to no staining of the negative control (HeLa KO for HeLa cells and E0771 for MDA-MB-231 cells) was selected for further EV stainings (indicated with an arrow). A) Antibody titration of CD81 APC and CD81 FITC, CD63 APC, CD9 FITC calculated with HeLa-EVs. B) Antibody titration of CD81 APC, CD63 APC (with illustration of dot plots obtained with the same antibody dilutions on MDA-MB-231- and E0771-EVs), CD63 FITC, CD9 FITC, CD29 AF488, CD29 AF647, CD44 AF647, SSEA-4 APC, CD98 APC.

**Sup. Figure 3:**
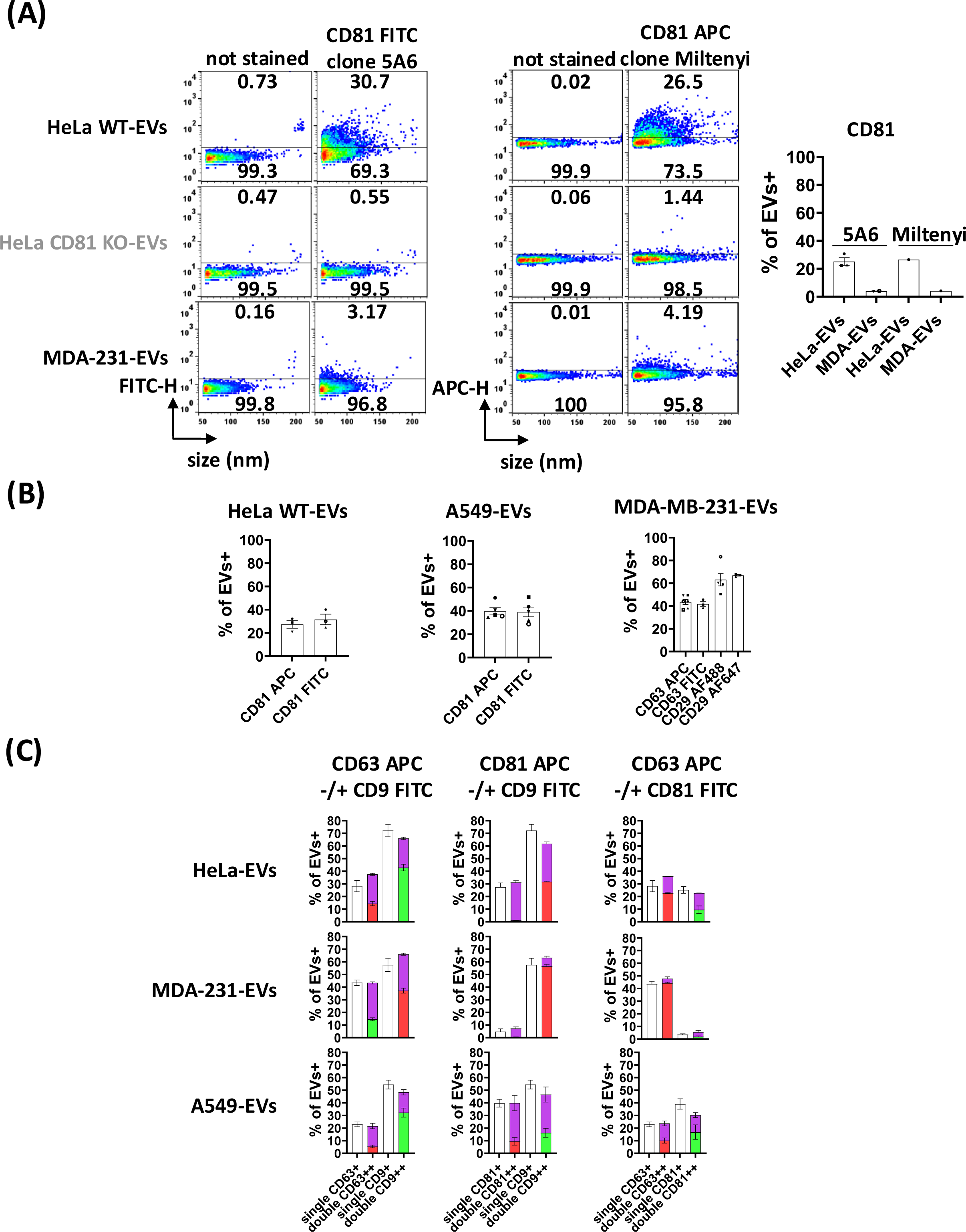
Efficacy of EV detection by different antibody clones, different fluorophores, and single-versus double-staining. A) CD81 detection by different antibody clones in HeLa- and MDA-MB-231-EVs (MDA-231-EVs). Dot plots of single-staining of HeLa WT-versus HeLa CD81 KO-EVs and MDA-MB-231-EVs for anti-CD81-FITC clone 5A6 and another anti-CD81-APC clone (Miltenyi antibody clone provided with the MacsPlexEV kit). The results are represented with size distribution in nm and fluorescence intensity (FITC-H or APC-H), and the percentages of EVs below and above the positivity threshold are indicated on the dot plots (left panel). Bar charts showing the percentage of labeled HeLa-EVs and MDA-MB-231-EVs for CD81-FITC clone 5A6 and for CD81-APC clone Miltenyi compared to the total population (mean ± SEM of n=1-3 independent experiments). B) Comparison of different fluorochromes. Hela WT-EVs (left) and A549-EVs (middle) labeled with anti-CD81 clone 5A6 coupled to either FITC or APC, and MDA-MB-231 EVs labeled with anti-CD63 coupled to either APC or FITC, or anti-CD29 coupled to either AF488 or AF647 (right) are shown (mean ± SEM of n=3-6 independent experiments). C) Comparison of the % of positive EVs calculated from single (white bars) versus various combinations of double-stainings (coloured bars), for each tetraspanine and each cell line as indicated. Colour code: green = % of single FITC-labeled EVs, red = % of single APC-labeled EVs, purple = % of double FITC/APC-labeled EVs. Note that all calculated percentages are comparable.

**Sup. Figure 4:**
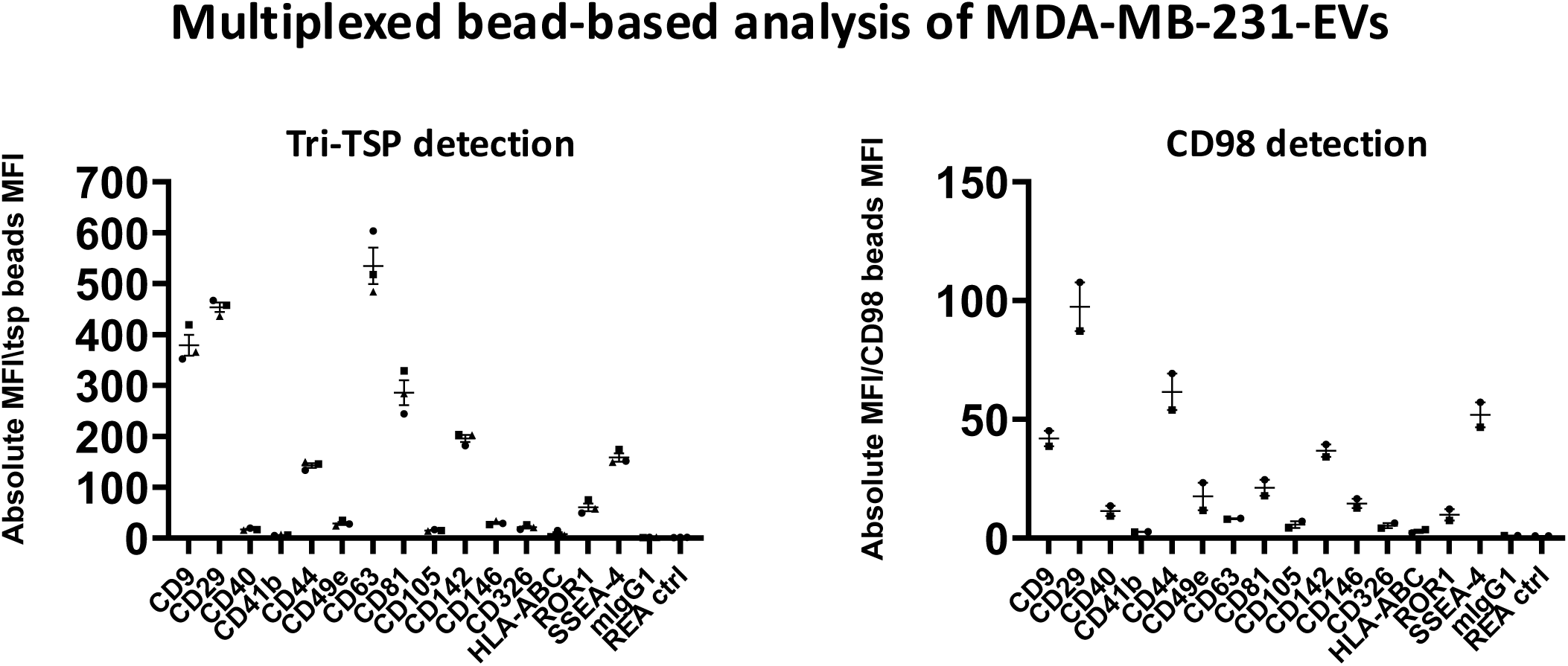
Analysis of other surface markers on MDA-MB-231 EVs by MacsPlexEV assay. EV surface proteins were profiled using a multiplex bead-based flow cytometry assay (MacsPlexEV, Miltenyi). Captured EVs were counterstained with APC-labeled detection antibodies (mixture of anti-CD9, anti-CD63 and anti-CD81 antibodies = Tri-TSP, left) or SLC3A2/CD98 (right). The 15 capture beads, out of the 37 tested, giving a positive signal compared to isotype controls are shown. CD41b, CD81, HLA-ABC and SSEA-4 were compared to REA control and the rest to IgG1 control. Results are presented as ratio of MFI (beads+EVs+detection antibodies) / MFI (beads+detection antibodies). In addition to the three tetraspanins, the strongest signals were observed for CD29/ITGB1, CD44, CD142, SSEA-4. We selected CD29, CD44 and SSEA-4 for further analysis by nano-flow cytometry.

**Sup. Figure 5:**
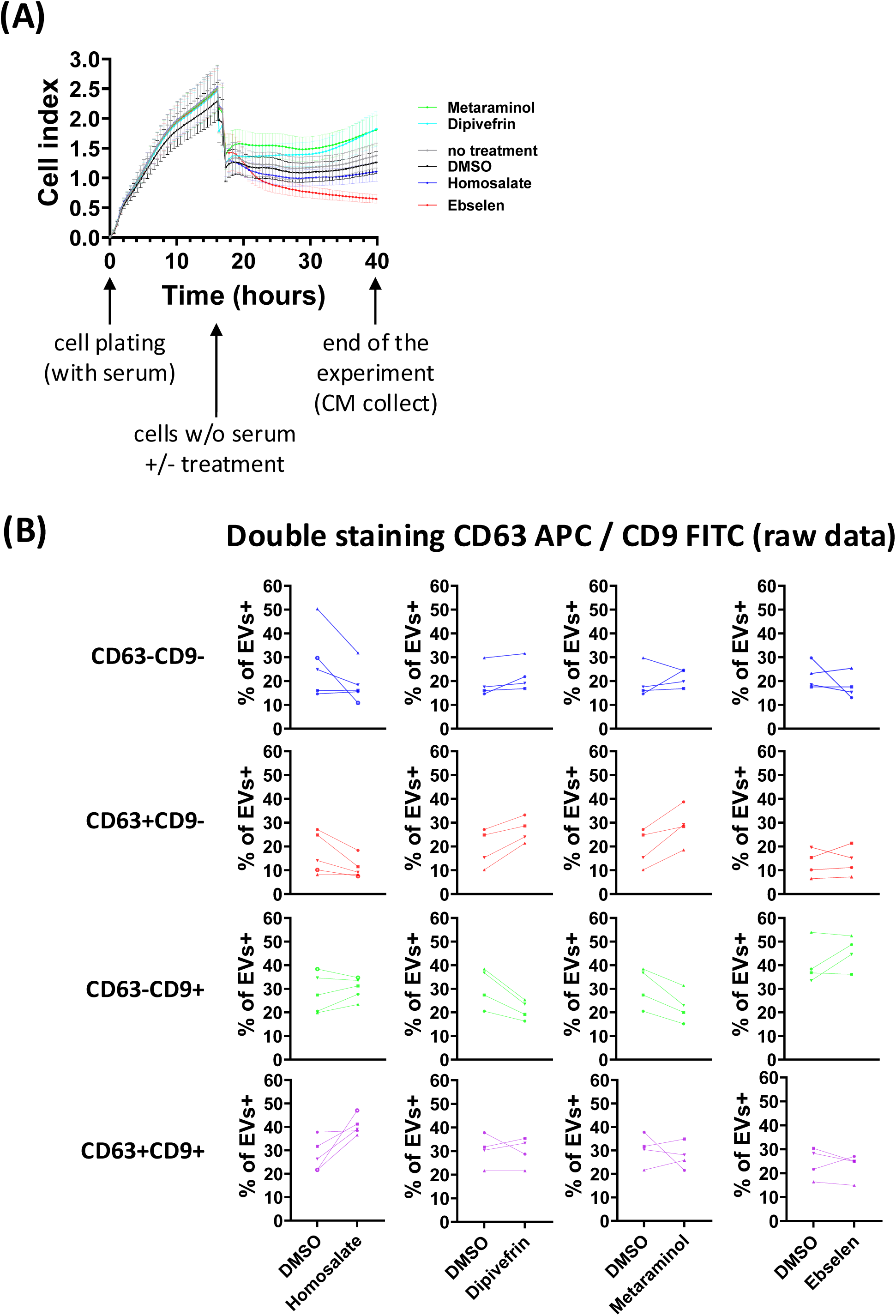
Effect of different pharmacological drugs on cell growth, and on the proportion of subtypes of EVs of MDA-MB-231 cells. A) MDA-MB-231 cells real-time evolution of cell index measured by xCELLIgence device (proportional to cell adhesion and number) before (0-16h) and after (16-40h) treatment with Homosalate, Dipivefrin, Metaraminol or Ebselen (10 μM) in absence of FCS, compared to DMSO as a control condition. Measurements were programmed every 30 min for a total time of 40 h (mean ±SEM of n=3 in duplicate). B) Raw data of the double stainings CD63 APC/CD9 FITC shown in Figure 4B of MDA-MB-231 EVs after drug treatment. Each graph shows the percentages of double-negative (blue), single-positive (red: APC or green: FITC) and double-positive (purple) EV populations released by non-treated (DMSO) and treated (Homosalate, Dipivefrin, Metaraminol, Ebselen) MDA-MB-231 cells in the same experiment.

**Sup. Table 1:** List of commercial antibodies used for the staining of EVs before Flow NanoAnalysis. Conc.: Concentration of commercial stock solution, Nb of Fluo/Ab: Number of fluorochromes per antibody, Opt conc.: Optimal concentration, n.d.: info not determined by provider.

|  | Company | Reference | Clone | Batch | Conc. (µg/ml) | Nb of Fluo/Ab | Opt conc. (µg/ml) |
| --- | --- | --- | --- | --- | --- | --- | --- |
| CD63 APC | Biolegend | 353008 | H5C6 | B373947 | 200 | 1,26 | 4 |
| CD63 FITC | Biolegend | 353006 | H5C6 | B262101 | 200 | 6,1 | 2 |
| CD81 APC | Biolegend | 349510 | 5A6 | B296062 | 200 | 1.10-1.19 | 4 |
| CD81 FITC | Biolegend | 349504 | 5A6 | B342883 | 200 | 6,46 | 4 |
| CD81 APC | Miltenyi | 130-108-813 | n.d. | n.d. | n.d. | n.d. | n.d. |
| CD9 FITC | Biolegend | 312104 | H19a | B359149 | 100 | 6,76 | 2 |
| CD98 APC | Miltenyi | 130-127-305 | REA387 | 5230904572 | 250 | n.d. | 5 |
| CD29 AF647 | Biolegend | 303017 | TS2/16 | B330207 | 400 | 3,27 | 2 |
| CD29 AF488 | Biolegend | 303015 | TS2/16 | B392842 | 200 | 4,22 | 2 |
| CD44 AF647 | Biolegend | 397511 | C44Mab-5 | B394673 | 200 | 5,43 | 4 |
| SSEA-4 APC | Biolegend | 330417 | MC-813-70 | B388834 | 50 | 1,45 | 1 |

**Sup. Figure 6:**
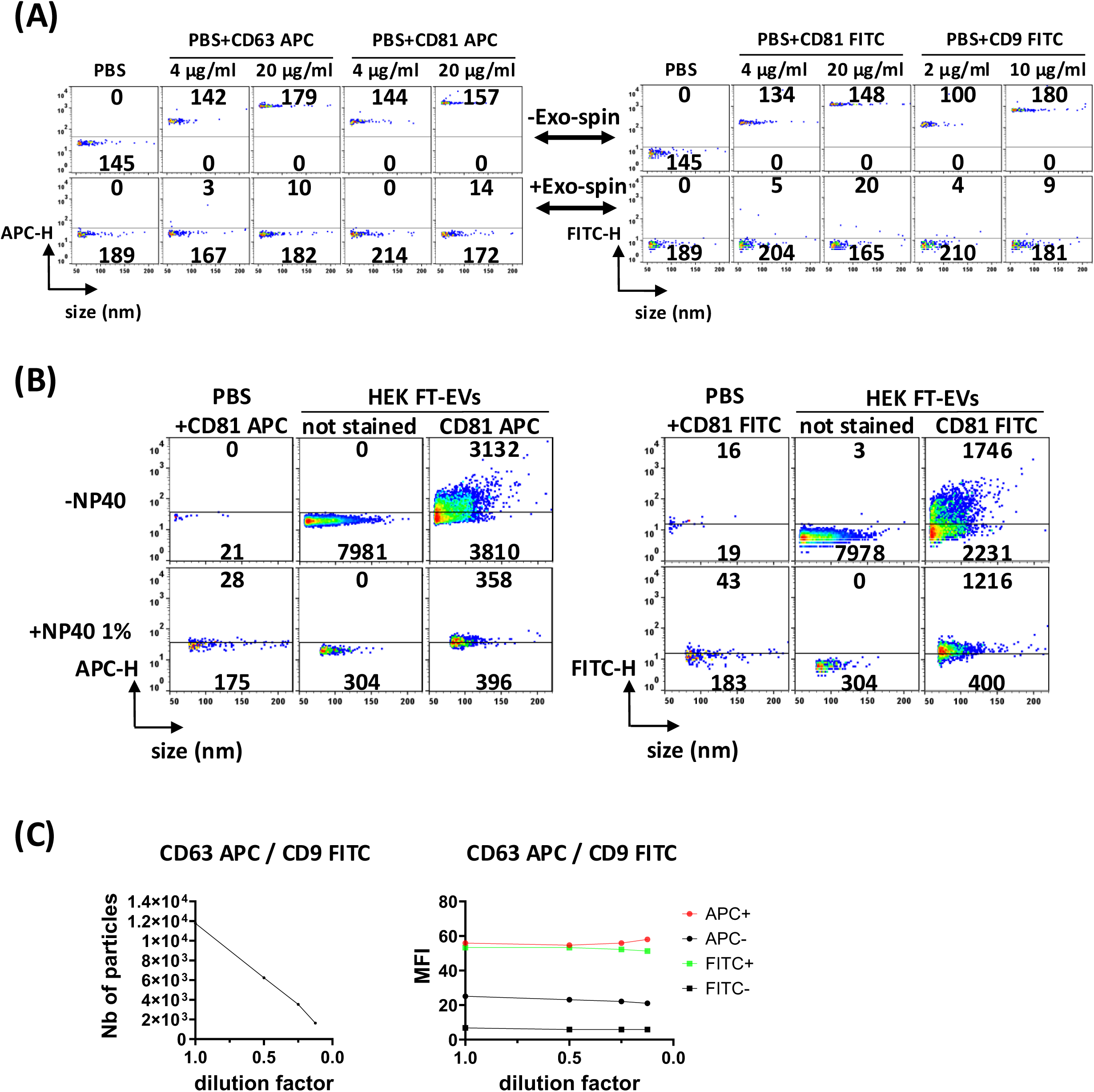
Control experiments for validation of labeled EV detection by Flow NanoAnalyzer. A) Dot plots of antibodies diluted in PBS (1 µl antibody pure or diluted 5X + 9 µl PBS) after 30 min incubation at 4°C and then diluted 18X with PBS before analysis by Flow NanoAnalyzer for the condition “-Exo-spin” (without SEC cleaning) or eluted with 180 µl of PBS for the condition “+Exo-spin” (with SEC cleaning). Results are represented with size distribution in nm and fluorescence intensity (APC-H or FITC-H), and the number of recorded events is indicated on each dot plot. Fluorescent antibody aggregates detected by the Flow NanoAnalyzer are eliminated by the SEC cleaning step. B) Dot plots of single staining of HEK293 FT-EVs for CD81 (APC or FITC) with their respective controls (antibody at the same concentration without EVs = PBS + CD81 APC/FITC, and EVs without antibody = not stained). Samples were analyzed first without detergent (top panels), and then treated with detergent (NP-40 1%) and analyzed immediately with the same settings by the Flow NanoAnalyzer (bottom panels): 89,1% (CD81-APC) or 59,4% (CD81-FITC) of particles are eliminated by NP40 treatment of EVs, with, however, the appearance of events/particles induced by NP40 treatment of the antibodies without EVs, representing 27% (CD81-APC) or 14% (CD81-FITC) of the events counted in the NP40-treated EV samples. C) Number of particles counted by Flow NanoAnalyzer upon serial dilution of the MDA-MB-231-EVs labeled with CD63-APC and CD9-FITC shows perfect correlation (left panel), and MFI is identical over all dilutions (right panel), thus excluding simultaneous detection of multiple EVs as single events (i.e. absence of swarm effect).

**Sup. Figure 7:**
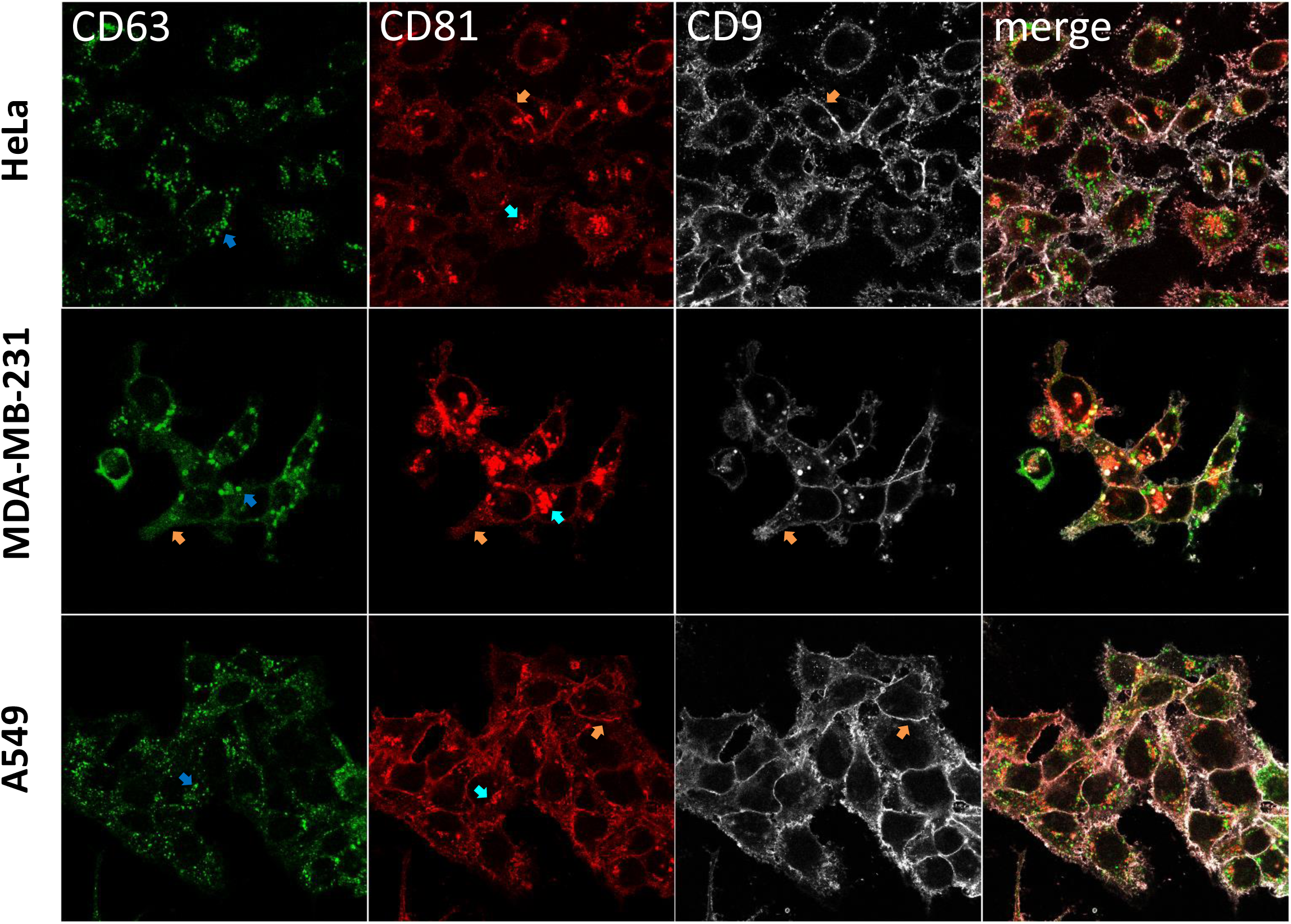
Immunofluorescence analysis of cellular location of CD63, CD81 and CD9 in the 3 cell lines. HeLa, MDA-MB-231 and A549 cells were fixed and stained for CD63 (green), CD81 (red) and CD9 (gray) in the presence of 0.1% saponin. Images were acquired on a Zeiss LSM 780 confocal microscope, one section in the middle of cells is shown. Orange arrows highlight examples of the plasma membrane with CD9 or CD81 staining in all 3 cell lines, and of CD63 at the plasma membrane only in MDA-MB-231. Blue arrows highlight examples of stained internal compartments: CD81 (light blue) or CD63 (dark blue) are both present in internal compartments, but generally not colocalized. CD9 is found in rare CD81 internal compartments in MDA-MB-231, but not in other cells. Scale bar: 10 µm.

**Sup. Table 2: MIFlowCyt-EV reporting framework.**

Model of excel spreadsheet proposed by MIFlowCyt-EV (Welsh *et al*., 2020) and MIFlowCyt, downloaded from https://www.evflowcytometry.org/resources/ and filled in to report specific settings of the experiments described in this article.

## REFERENCES

Brealey J, Lees R, Tempest R, Law A, Guarnerio S, Maani R, Puvanenthiran S, Peake N, Pink R, Peacock B (2024) Shining a light on fluorescent EV dyes: Evaluating efficacy, specificity and suitability by nano-flow cytometry. J Extracell Biol 3: e70006

Buzas EI (2023) The roles of extracellular vesicles in the immune system. Nat Rev Immunol 23: 236–250

Chen C, Cai N, Niu Q, Tian Y, Hu Y, Yan X (2023) Quantitative assessment of lipophilic membrane dye-based labelling of extracellular vesicles by nano-flow cytometry. J Extracell Vesicles 12: e12351

Cocozza F, Grisard E, Martin-Jaular L, Mathieu M, Thery C (2020a) SnapShot: Extracellular Vesicles. Cell 182: 262–262 e261

Cocozza F, Nevo N, Piovesana E, Lahaye X, Buchrieser J, Schwartz O, Manel N, Tkach M, Thery C, Martin-Jaular L (2020b) Extracellular vesicles containing ACE2 efficiently prevent infection by SARS-CoV-2 Spike protein-containing virus. J Extracell Vesicles 10: e12050

Daaboul GG, Gagni P, Benussi L, Bettotti P, Ciani M, Cretich M, Freedman DS, Ghidoni R, Ozkumur AY, Piotto C et al (2016) Digital Detection of Exosomes by Interferometric Imaging. Sci Rep 6: 37246

Dixson AC, Dawson TR, Di Vizio D, Weaver AM (2023) Context-specific regulation of extracellular vesicle biogenesis and cargo selection. Nat Rev Mol Cell Biol 24: 454–476

Fan SJ, Kroeger B, Marie PP, Bridges EM, Mason JD, McCormick K, Zois CE, Sheldon H, Khalid Alham N, Johnson E et al (2020) Glutamine deprivation alters the origin and function of cancer cell exosomes. EMBO J: e103009

Fordjour FK, Guo C, Ai Y, Daaboul GG, Gould SJ (2022) A shared, stochastic pathway mediates exosome protein budding along plasma and endosome membranes. J Biol Chem: 102394

Gorgens A, Bremer M, Ferrer-Tur R, Murke F, Tertel T, Horn PA, Thalmann S, Welsh JA, Probst C, Guerin C et al (2019) Optimisation of imaging flow cytometry for the analysis of single extracellular vesicles by using fluorescence-tagged vesicles as biological reference material. J Extracell Vesicles 8: 1587567

Grisard E, Nevo N, Lescure A, Doll S, Corbe M, Jouve M, Lavieu G, Joliot A, Nery ED, Martin-Jaular L et al (2022) Homosalate boosts the release of tumour-derived extracellular vesicles with protection against anchorage-loss property. J Extracell Vesicles 11: e12242

Holcar M, Maric I, Tertel T, Goricar K, Cegovnik Primozic U, Cerne D, Giebel B, Lenassi M (2025) Comprehensive Phenotyping of Extracellular Vesicles in Plasma of Healthy Humans - Insights Into Cellular Origin and Biological Variation. J Extracell Vesicles 14: e70039

Lannigan J, Erdbruegger U (2017) Imaging flow cytometry for the characterization of extracellular vesicles. Methods 112: 55–67

Lees R, Tempest R, Law A, Aubert D, Davies OG, Williams S, Peake N, Peacock B (2022) Single Extracellular Vesicle Transmembrane Protein Characterization by Nano-Flow Cytometry. J Vis Exp

Loconte L, Arguedas D, El R, Zhou A, Chipont A, Guyonnet L, Guerin C, Piovesana E, Vazquez-Ibar JL, Joliot A et al (2023) Detection of the interactions of tumour derived extracellular vesicles with immune cells is dependent on EV-labelling methods. J Extracell Vesicles 12: e12384

Martin-Jaular L, Nevo N, Schessner JP, Tkach M, Jouve M, Dingli F, Loew D, Witwer KW, Ostrowski M, Borner GHH et al (2021) Unbiased proteomic profiling of host cell extracellular vesicle composition and dynamics upon HIV-1 infection. EMBO J 40: e105492

Mathieu M, Nevo N, Jouve M, Valenzuela JI, Maurin M, Verweij FJ, Palmulli R, Lankar D, Dingli F, Loew D et al (2021) Specificities of exosome versus small ectosome secretion revealed by live intracellular tracking of CD63 and CD9. Nat Commun 12: 4389

Morales-Kastresana A, Telford B, Musich TA, McKinnon K, Clayborne C, Braig Z, Rosner A, Demberg T, Watson DC, Karpova TS et al (2017) Labeling Extracellular Vesicles for Nanoscale Flow Cytometry. Sci Rep 7: 1878

Nolte-’t Hoen EN, van der Vlist EJ, Aalberts M, Mertens HC, Jan Bosch B, Bartelink W, Mastrobattista E, van Gaal EV, Stoorvogel W, Arkesteijn GJ et al (2012) Quantitative and qualitative flow cytometric analysis of nano-sized cell-derived membrane vesicles. Nanomedicine 8: 712–720

Palmulli R, Couty M, Piontek MC, Ponnaiah M, Dingli F, Verweij FJ, Charrin S, Tantucci M, Sasidharan S, Rubinstein E et al (2024) CD63 sorts cholesterol into endosomes for storage and distribution via exosomes. Nat Cell Biol 26: 1093–1109

Ricklefs FL, Maire CL, Reimer R, Duhrsen L, Kolbe K, Holz M, Schneider E, Rissiek A, Babayan A, Hille C et al (2019) Imaging flow cytometry facilitates multiparametric characterization of extracellular vesicles in malignant brain tumours. J Extracell Vesicles 8: 1588555

Rous BA, Reaves BJ, Ihrke G, Briggs JA, Gray SR, Stephens DJ, Banting G, Luzio JP (2002) Role of adaptor complex AP-3 in targeting wild-type and mutated CD63 to lysosomes. Mol Biol Cell 13: 1071–1082

Roux Q, Van Deun J, Dedeyne S, Hendrix A (2020) The EV-TRACK summary add-on: integration of experimental information in databases to ensure comprehensive interpretation of biological knowledge on extracellular vesicles. J Extracell Vesicles 9: 1699367

Saftics A, Abuelreich S, Romano E, Ghaeli I, Jiang N, Spanos M, Lennon KM, Singh G, Das S, Van Keuren-Jensen K et al (2023) Single Extracellular VEsicle Nanoscopy. J Extracell Vesicles 12: e12346

Schurz M, Danmayr J, Jaritsch M, Klinglmayr E, Benirschke HM, Matea CT, Zimmerebner P, Rauter J, Wolf M, Gomes FG et al (2022) EVAnalyzer: High content imaging for rigorous characterisation of single extracellular vesicles using standard laboratory equipment and a new open-source ImageJ/Fiji plugin. J Extracell Vesicles 11: e12282

Spitzberg JD, Ferguson S, Yang KS, Peterson HM, Carlson JCT, Weissleder R (2023) Multiplexed analysis of EV reveals specific biomarker composition with diagnostic impact. Nat Commun 14: 1239

Susa KJ, Seegar TC, Blacklow SC, Kruse AC (2020) A dynamic interaction between CD19 and the tetraspanin CD81 controls B cell co-receptor trafficking. Elife 9

Tian Y, Gong M, Hu Y, Liu H, Zhang W, Zhang M, Hu X, Aubert D, Zhu S, Wu L et al (2020) Quality and efficiency assessment of six extracellular vesicle isolation methods by nano-flow cytometry. J Extracell Vesicles 9: 1697028

Tian Y, Ma L, Gong M, Su G, Zhu S, Zhang W, Wang S, Li Z, Chen C, Li L et al (2018) Protein Profiling and Sizing of Extracellular Vesicles from Colorectal Cancer Patients via Flow Cytometry. ACS Nano 12: 671–680

Tkach M, Thalmensi J, Timperi E, Gueguen P, Nevo N, Grisard E, Sirven P, Cocozza F, Gouronnec A, Martin-Jaular L et al (2022) Extracellular vesicles from triple negative breast cancer promote pro-inflammatory macrophages associated with better clinical outcome. Proc Natl Acad Sci U S A 119: e2107394119

van Niel G, Carter DRF, Clayton A, Lambert DW, Raposo G, Vader P (2022) Challenges and directions in studying cell-cell communication by extracellular vesicles. Nat Rev Mol Cell Biol 23: 369–382

Welsh JA, Van Der Pol E, Arkesteijn GJA, Bremer M, Brisson A, Coumans F, Dignat-George F, Duggan E, Ghiran I, Giebel B et al (2020) MIFlowCyt-EV: a framework for standardized reporting of extracellular vesicle flow cytometry experiments. J Extracell Vesicles 9: 1713526

Yates AG, Pink RC, Erdbrugger U, Siljander PR, Dellar ER, Pantazi P, Akbar N, Cooke WR, Vatish M, Dias-Neto E et al (2022a) In sickness and in health: The functional role of extracellular vesicles in physiology and pathology in vivo: Part I: Health and Normal Physiology: Part I: Health and Normal Physiology. J Extracell Vesicles 11: e12151

Yates AG, Pink RC, Erdbrugger U, Siljander PR, Dellar ER, Pantazi P, Akbar N, Cooke WR, Vatish M, Dias-Neto E et al (2022b) In sickness and in health: The functional role of extracellular vesicles in physiology and pathology in vivo: Part II: Pathology: Part II: Pathology. J Extracell Vesicles 11: e12190

Zhu S, Ma L, Wang S, Chen C, Zhang W, Yang L, Hang W, Nolan JP, Wu L, Yan X (2014) Light-scattering detection below the level of single fluorescent molecules for high-resolution characterization of functional nanoparticles. ACS Nano 8: 10998–11006

